# An Integrated Proteomics and Genomics Approach to Identify Essential Protein Kinases During Human Trophoblast Development

**DOI:** 10.64898/2026.07.23.740390

**Authors:** Rajnish Kumar, Purbasa Dasgupta, Soma Ray, Soumen Paul

## Abstract

In the developing human placenta, three subtypes of trophoblast cells, cytotrophoblasts (CTBs), extravillous trophoblasts (EVTs), and syncytiotrophoblasts (STBs), mediate critical functions essential for a successful pregnancy. CTBs constitute the stem/progenitor compartment and differentiate into STBs and EVTs within the floating and anchoring villi, respectively. STBs establish the maternal-fetal exchange interface and secrets human chorionic gonadotropin (hCG), a hormone vital for the maintenance of early pregnancy. EVTs anchor the maternal endometrium and invade the uterine tissue to remodel maternal cells, supporting implantation and progression of pregnancy. In this study, we used human trophoblast stem cells (hTSCs) as a model system and performed quantitative, label-free liquid chromatography–tandem Mass Spectrometry (LC-MS/MS) to profile proteome and phosphoproteome in TSC stem state (analogous to undifferentiated CTBs) and following their differentiation to STBs and EVTs. Through a multiomics approach, we integrated our proteomics data with global gene expression profiles to correlate cell-type specific gene and protein expression during human trophoblast development. We also identified global phosphoproteome and analyzed kinases that are specifically active in hTSC stem state, as well as in differentiated STBs and EVTs. We experimentally validated specific kinases, such as BUB1B, PAK6, PKYMT1 and TNIK are essential for maintaining the hTSC stem-state. Additionally, atypical protein kinase C isoforms PKCζ is essential for STB development, while PTK2B, SRC, TRIO and LYN are important for EVT development. Our findings highlight key kinases uniquely required for specific stages of trophoblast development during human placentation and suggest that pharmacological inhibition of these kinases could negatively impact the placentation process during pregnancy.

## Introduction

Normal development of the placenta ensures successful pregnancy (1, 2). Although placenta is a transient organ, it mediates diverse functions including supplying of oxygen and nutrient to the developing fetus as well as endocrine functions that facilitate maternal adaptation for continuation of pregnancy (1–3). Most of these placental functions are mediated by the trophoblast cell lineage.

After implantation of a human blastocyst, a primitive syncytium supports the initial embryonic development (4, 5). Eventually, the primitive syncytium is replaced by a villous placenta, which supports the pregnancy to term. In a developing villous human placenta, the CTBs establish the stem/progenitor cell compartment (4–6). At the onset of placenta development CTBs undergo extensive proliferation that ensures growth of the placenta. CTBs adapt two differentiation pathways in two different types of placental villi (4, 5). In floating villi, which floats into the maternal blood the CTBs differentiate to multinucleated STBs. Thus, the multinucleated STB of a human placenta is in direct contact with the maternal blood and functions as the interface of gas and nutrient exchange. The STB layer also provides the barrier function to protect the fetus from infectious and toxic agents. In anchoring villi, which anchors the placenta to the maternal endometrium, CTBs adopt an invasive differentiation fate and develop into invasive EVTs (7–9). Whereas CTB to STB development is associated with arrested replication and extensive cell-fusion, the CTB to EVT differentiation is initially associated with extensive replication along with alteration of gene expression program to establish a CTB cell column, known as column CTBs. Eventually the column CTBs at the distal end of the cell column acquire invasive phenotype to develop into two types of EVTs; (i) interstitial EVTs, which remains in the uterine interstitium and module the function of uterine cells via endocrine functions, and (ii) endovascular EVTs, which infiltrate and remodel the uterine artery (7–9) to create dilated, low resistance blood vessels (8, 10) required to meet the nutrient demands of the developing fetus. Thus, three trophoblast subtypes of a developing human placenta are characterized with distinct biological processes, such as self-renewal vs. differentiation balance in CTBs, cell fusion and transport function in the STBs and invasion, vascular remodeling and endocrine function in EVTs.

Numerous transcriptomics, genomics and epigenomics studies have provided important insights into the gene regulatory dynamics during human trophoblast development (11–20). A global proteomics analyses was conducted with second trimester human placentae to identify proteome in invading CTBs (21). While developing this manuscript, another study (22) reported dynamic protein expression pattern in trophoblast cells using preimplantation human embryos and hTSC differentiation model. However, an integrative approach to correlate gene expression with the proteome and phosphoproteome to understand the dynamics of signaling pathways in different human trophoblast subtypes has not yet been undertaken. An earlier study (23) performed integrative analysis of proteome, phosphoproteome and transcriptome in two development stages (Gestational day 7.5 and 9.5) during mouse placentation. However, trophoblast development in mouse versus humans is associated with fundamental differences. Therefore, in this study, using the label free LC-MS/MS approach (24, 25) we have performed both global proteomics and phosphoproteomics with hTSCs in their stem state and after they are differentiated to STBs and EVTs. We integrated the global proteomics data with global gene expression data (RNA-seq) in hTSCs along and single-cell resolution gene expression (single-cell RNA-seq) data in first-trimester human placentae to correlate biological pathways that are associated with distinct stages of trophoblast development. Through integrated analyses of expression patterns and activity, we identified active protein kinases in undifferentiated hTSCs and upon induction of STB and EVT fates. Finally, we used specific inhibitors to identify which of these kinases are essential to maintain hTSC stem state and for their differentiation to STBs and EVTs. Our study identifies expression of unique regulatory proteins, including specific kinases, which are essential for different stages of trophoblast development during human placentation and could be an important resource for future studies to better understand normal and pathological human pregnancies.

## Materials and Methods

### Human trophoblast stem cells culture

CT27 hTSC line was kindly shared by Dr. Hiroaki Okae (Tohoku University, Japan). The cells were maintained, passaged, and differentiated as reported in (11, 26). The cells were tested for mycoplasma using mycoplasma detection kit (ATCC, 30-1012K). Culture conditions are briefly mentioned below. All associated reagents and their sources to culture hTSCs in stem state and STB/EVT differentiation are mentioned in Supplementary Table 1.

#### hTSC culture proliferative condition

Cell were maintained in hTSC stem-state culture medium comprised of DMEM/F12 supplemented with 0.1mM 2-mercaptoethanol, 0.2% FBS, 0.5% Penicillin-Streptomycin, 0.3% BSA, 1% ITS- X supplement, 1.5μg/ml L-ascorbic acid, 50ng/ml EGF, 2μM CHIR99021, 0.5μM A83-01, 1μM SB431542, 0.8mM VPA and 5μM Y27632. Cells were cultured at 37°C in 5% CO2 and the culture media was replaced every 2 days. When the cells reached 60% to 80% confluence, they were dissociated using TrypLE for 5-7 minutes at 37°C and passaged to a new Collagen IV coated plate.

#### EVT differentiating culture condition

For EVT differentiation, hTSCs (1×10^5^ cells in each well of a 6 well tissue culture plate) were seeded in EVT culture medium, composed of DMEM/F12 supplemented with 0.1mM 2-mercaptoethanol, 0.5% Penicillin-Streptomycin, 0.3% BSA, 1% ITS-X supplement, 100 ng/ml NRG1, 7.5 μM A83-01, 2.5μM Y27632 and 4% KnockOut Serum Replacement. Matrigel was added to the medium at a final concentration of 2% after the cells were resuspended in the media. On day 3, the medium was replaced with EVT medium without NRG1 and the Matrigel was added to a final concentration of 0.5%. Cells were cultured in this condition for another 3 days, when most of the cells undergo EVT differentiation.

#### STB differentiating culture condition

For the induction of 2D culture of STB from hTSCs, cells were seeded in 6-well plate (1×10^5^ cells in each well) coated with 2.5μg/ml Collagen IV in a density of 1 × 10^5^ cells per well and cultured with STB medium, which is composed of DMEM/F12 supplemented with 0.1mM 2-mercaptoethanol, 0.5% Penicillin-Streptomycin, 0.3% BSA, 1% ITS- X supplement, 2.5μM Y27632, 4% forskolin and 4% KnockOut Serum Replacement. The medium was replaced after 3 days, and the cells were analyzed on day 6.

### Preparation of protein samples for proteomics analyses

For proteomics ananlyses 4×10^6^ hTSCs in stem state or after differentiation to STBs and EVTs were lysed by resuspending in 200ul of RIPA buffer with protease, phosphatase inhibitors and nuclease and incubated on ice for 30 minutes followed by sonication. For each condition 4 sample sets were used. Samples were then centrifuged at 14000g for 10 minutes at 4^0^C and supernatant were transferred to new tubes. Samples were dried in a speedvac and resuspended in 100ul of 50mM Triethylammonium bicarbonate. Disulfides were reduced by adding 10ul of 50mM Tris(2-carboxyethyl) phosphine (5mM final) and incubating at 55^0^C for 30 minutes. Cysteines were alkylated by adding 3ul of 375mM Iodoacetamide (10mM final, I5161, Sigma, St. Louis, MO, USA) and incubating at RT in the dark for 30 minutes. Proteins were precipitated by adding 450ul of acetone (1:5 dilution, 270725, Sigma) and incubating at -20^0^C overnight. Samples are centrifuged at 14000g for 10 minutes at 4^0^C. Supernatant was discarded while allowing the proteins to air dry on the bench top for 15 minutes. Proteins were resuspended in 100ul of 50mM Triethylammonium bicarbonate buffer (T7408 Sigma), 2mM CaCl2 and digested by adding 500ng of trypsin (0.1 mg/mL, T6567, Sigma) and incubating overnight at 37^0^C at 500 RPM. Reactions were quenched by the addition of 10% formic acid (5.33002, Sigma) to 1%. Peptides were serially enriched using the TiO2 (A32993, Thermo-Fisher, Waltham, MA, USA) followed by the Fe-NTA enrichment kits (A32992, Thermo-Fisher) and the flow through from the FeNTA kit was used for global proteomic samples. Peptide concentrations were determined using the nanodrop UV system and based on the concentrations, 10ul of each sample was loaded onto the C18 RP column (059143, Thermo-Fisher) and injected into the Orbitrap Ascend mass spectrometer with FAIMS for LC-MS/MS analysis.

### Analysis of Global Proteomics Data

Peptide identification and quantification were done using the Proteome Discoverer 3.0, in which SeQuest software was used against the human proteomics database downloaded from Uniprot database on 05-05-2023. The files with identified proteins were imported in RStudio environment. We have used DEP2 v 0.5.28.2 tool (27) for processing proteins containing files. In the first step make_unique() function was used to make data set unique which was further provided into make_se() function to construct SummarizedExperiement object. The normalization of the data was done using normalize_vcn() function. In the next step imputation was done using impute() function with “MinProb” as fun argumental parameter. The differential profiling was done using test_diff() function and result was stored into SummarizedExperiement object using add_rejection() function. The sample-to-sample correlation heatmap was plotted using plot_cor() function. Gene set enrichment analysis was performed using test_GSEA() function of DEP2 where we provided “Human” and “GO” as argument parameters for species & type respectively.

### Analysis of Phosphoproteomics Data

For phosphoproteomics analyses Proteome Discover 3.0 tool was used for identification and quantification. Data was searched using the Proteome Discoverer 3.0 SeQuest software against the human proteome database. Quantification & identification of phospho proteins & peptide files were imported into RStudio workspace. We have used DEP2 tool for downstream analysis following similar steps mentioned for global proteomics analysis above. We wrote a Python script to subset peptides containing phosphorylation events. In the next step we have quantified the types of phosphorylation events, such as mono, di or three & more than three phosphorylation events. Also, we have quantified the distribution of the phosphorylated Serine (S), Threonine (T) and Tyrosine (Y) residues. For identifying kinases in different cell types, we downloaded the reported human kinome from the KinHub database. Next using Python script, we subset the differentially expressed kinases from phosphoproteome of three trophoblast cell types. Finally, we used RokaiXplorer computational tool to identify phosphokinase activity.

### Single-Cell RNA Sequencing and Analysis

Details of single-cell RNA-seq analyses with first-trimester human placenta were reported earlier (28). Briefly, single-cell suspensions from two first-trimester placentae were generated and transcriptomic profiles were obtained using the 10x Genomics Chromium Single Cell Gene Expression Solution (10xgenomics.com). The 10x Genomics Loupe Cell Browser software was used to find significant genes, cell types, and substructure within the single-cell data.

### RNA-Seq Analyses

The RNA-seq datasets used in this study were generated in the Paul laboratory from wild-type CT27 hTSCs cultured under stem-state conditions or differentiated into STBs and EVTs. Although all datasets were generated simultaneously, the STB RNA-seq dataset was previously analyzed in an earlier publication (17) and raw data was deposited to NCBI GEO database under accession GSE214634. The hTSC and EVT RNA-seq datasets generated from the same experiment are deposited under GEO accession GSE296084. The RNA-seq analysis was performed according to published protocol (17, 26). Total RNA from hTSCs, cultured either in stem state or upon STB and EVT differentiation were isolated using RNeasy Mini Kit (74104 Qiagen, Germantown, MD, USA) following manufacturer’s protocol. Integrity of the total RNA samples was evaluated using an Agilent Technologies 2100 Bioanalyzer. The total RNA fraction was processed by oligo dT bead capture of mRNA, fragmentation, and reverse transcription into cDNA. After ligation with the appropriate Unique Dual Index (UDI) adaptors, the cDNA library was prepared using the Universal Plus mRNA-seq +UDI library preparation kit (NuGEN; Marion, SD). Adaptor and low-quality reads were removed using Rfastp R package. For the current study, RNA-seq reads from all three cell types were processed using an identical analysis pipeline. Reads were aligned to the human reference genome (GRCh38, Ensembl Release 92) using STAR v2.6.1c, transcript abundance and raw gene counts were quantified using RSEM v1.3.1, and differential gene expression analyses were performed using DESeq2 (Bioconductor).

### Integration of RNA-seq and Global Proteomics Data

We integrated the unique set of proteins, which are expressed in hTSCs, STBs and EVTs and expression of corresponding genes in trophoblast cells by integrating global proteomics data with the global RNA-seq data. We wrote a Python script, which utilizes the concept of dictionary to hold the key and its value. For each of the differential conditions, we have created one dictionary for RNA-Seq where we stored Gene name as key and log2(Fold change) as values. Similarly, for the same differential condition in Proteomics data, we have created a dictionary for proteins where we put the gene corresponding to the proteins as key and log2(Fold change) as value.

The example is given below

Gene_dict = {“Gene1”: “Log 2(Fold change) of Gene1”, “Gene2”: “Log 2(Fold change) of Gene2”,..”, “GeneN”: “Log 2(Fold change) of GeneN”}, where N is the number of differentially expressed genes.

Prot_dict= {“Gene1of Protein1”: “Log 2(Fold change) Protein1”, “Gene1of Protein2”: “Log 2(Fold change) Protein2”,..,“Gene1of ProteinM”: “Log 2(Fold change) ProteinM “}, where M is number of differentially expressed proteins.

Now using the common genes, a new dictionary was created, where values were stored as a list corresponding to the gene name as a key in above mentioned dictionaries. We followed the following dictionary structure.

Common_dictionary = {“Gene1”: “[‘log2(Fold change of RNAseq)’, ‘log2(Fold change of corresponding protein 1 from Proteomics)’]”, {“Gene2”: “[‘log2(Fold change of RNAseq)’, ‘log2(Fold change of corresponding protein 2 from Proteomics)’]”, ..-, {“GeneK”: “[‘log2(Fold change of RNAseq)’, ‘log2(Fold change of corresponding protein 2 from Proteomics)’]”}, where K is number of genes found in both differential RNA-Seq and Proteomics data.

Based on the sign of Fold change value of Proteomics and RNA-Seq data, the combined data was separated into files containing correlated or anticorrelated genes and proteins data. The correlated dataset was taken into RStudio workspace and gene set enrichment was performed using clusterProfiler Bioconductor package for each of the differential conditions. Volcano plot generated for each of the differential condition using ggplot2 R package.

### Identification of Common and Exclusive Cell-type Proteins and Pathways

The result files from Proteome Discoverer 3.0 were imported in RStudio workspace. We have filter out the proteins based on their expression in at least 2 samples out of 4 for both global & phosphoproteomics data. Gene set enrichment analysis (GSEA) and Metascape web tools were used for gene ontology analyses of identified proteins to define biological functions, cellular processes, and cellular components.

### MTT Assay

For determining working concentration of kinase inhibitors, MTT Assay was performed using a commercially available kit (ab21109, Abcam, Waltham, MA, USA) following the manufacturer’s protocol. 3000 cells/well were seeded in 96-well plates (92097, Sigma) with various inhibitor concentrations. DMSO was used as control. Media change with inhibitor was given on day 3. On Day 4 (at 96 hours), 50ul of MTT reagent was added and cells were incubated at 37 °C for 3 hours in the dark to allow reduction of the tetrazolium salt to formazan by metabolically active cells. Formazan crystals were solubilized with provided solvent and absorbance was measured at 590 nm using a microplate reader. Cell viability was normalized to DMSO controls. Experiments were performed with three independent biological replicates.

### Studies with Kinase Inhibitors

Final experimental concentrations for kinase inhibitors were determined based on MTT assay in hTSC stem state and mentioned in Table 1. For cell proliferation assays, equal numbers of cells (4 × 10^4^) were seeded in each well of a 6-well plate and treated with either drug or DMSO (vehicle control) at the selected concentration. Cell growth was monitored over three consecutive days, after which cells were counted and processed for downstream analyses. For EVT and STB differentiation assays, 1 × 10^5^ drug-treated hTSCs were seeded under EVT or STB-specific differentiation conditions using the respective culture media in a 6-well plate. Cellular morphology was documented on day 6 of EVT differentiation and on day 5 of ST differentiation, followed by RNA and protein analyses of drug-treated cells.

**Table 1:**
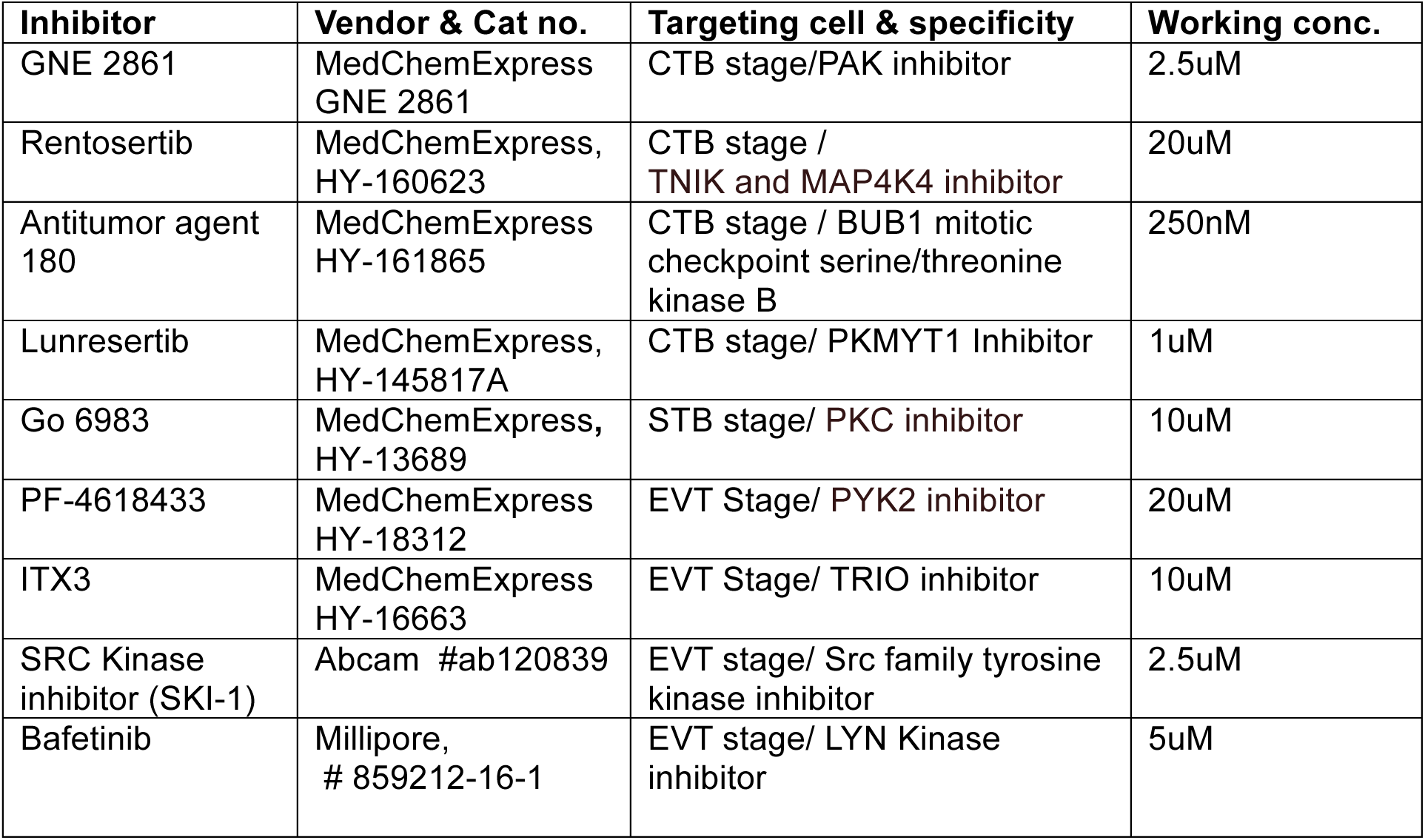
Kinase Inhibitors Used For the Study.

### Annexin-V/FITC Assay

Apoptosis of hTSCs in presence of kinase inhibitors was assessed using the Annexin V Apoptosis Detection Kit (BMSF00FI, Invitrogen, Carlsbad, CA, USA) according to the manufacturer’s instructions. Briefly, 5 × 10^5^ cells were seeded on 10 cm plates and cultured with DMSO or kinase inhibitors (at concentrations mentioned in Table 1) for 3 days. On day 4, cells were washed once with PBS, and trypsinized. Cells were resuspended in 1X Annexin V binding buffer and were stained with fluorochrome-conjugated Annexin V and propidium iodide (PI) and incubated for 15 min at room temperature in the dark. Unstained cells (DMSO treated) were kept for control. Following incubation, additional binding buffer was added, and samples were analyzed immediately by flow cytometry.

Data were acquired using a flow cytometer (Aurora) and analyzed with SpectroFlo software. Early apoptotic cells were defined as Annexin V-positive and PI-negative, whereas late apoptotic/necrotic cells were Annexin V-positive and PI-positive. The percentage of apoptotic cells was quantified relative to total cell population.

### RNA isolation and RT-qPCR

RNA extraction was carried out using RNeasy mini kit (Qiagen, 74104) according to the manufacturer’s protocol. RNA quantification was carried out by Nanodrop (Invitrogen). 1ug of RNA was used for cDNA synthesis using random hexamer primer (Thermo Scientific S0142) and Reverse Transcriptase enzyme (Invitrogen, 28025-013) by following standard protocols. qPCR was performed with SYBR Green master mix (Applied Biosystems, 4367659) and was performed in biological triplicates and 18sRNA was used as a housekeeping gene for normalization. GraphPad Prism (version 11) was used for analysis and visualization. Primer sequences for RT-qPCR analyses are mentioned below.

### Primers Used for RT-qPCR

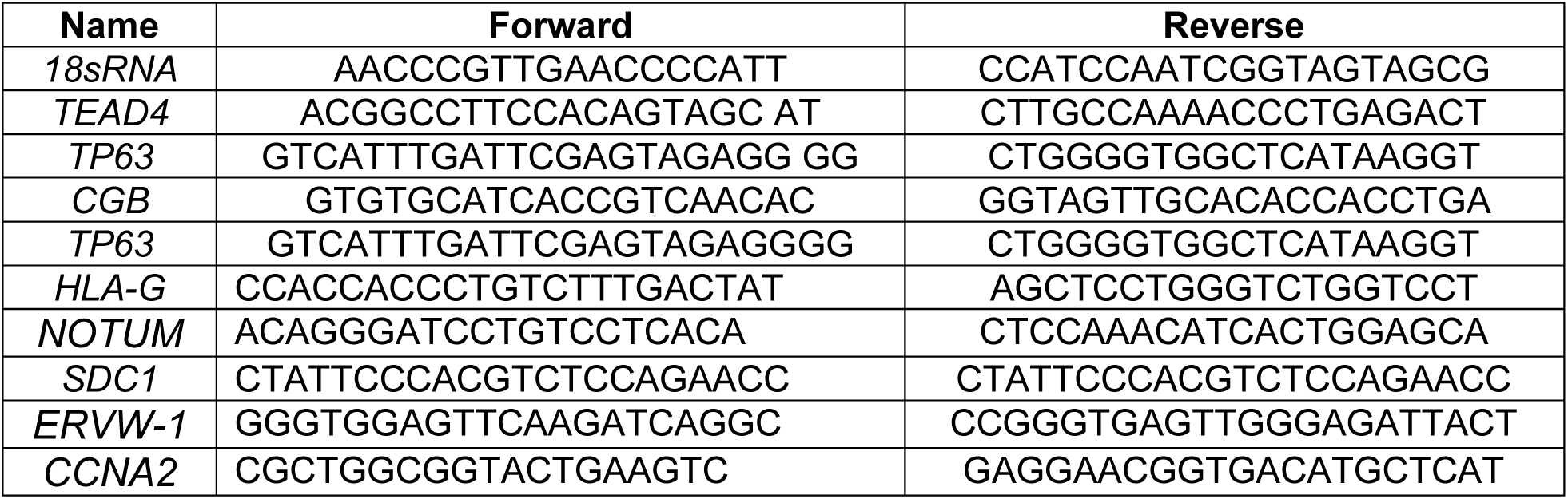

### Immunofluorescence

Cells were fixed with 4% paraformaldehyde/PBS at 4°C for 10 minutes, permeabilized with PBS-0.25%TritonX at room temperature for 15 minutes and blocked in PBS-0.1%TritonX-10%FBS at room temperature for an hour. Cells were incubated with primary antibodies in PBS-0.1%TritonX-10%FBS at 4°C overnight. On the next day, they were stained with secondary antibody in PBS-0.1%TritonX-10%FBS at room temperature for 2 hours. DAPI (564907, BD Biosciences, San Jose, CA, USA) counterstain was done and then antifade mounting media (Invitrogen, S36936) were used. Anti-CDH1 (Abcam 1416), anti-KI67 (Abcam, 16667), anti-HLA-G (Abcam 7758), and anti-hCG-β (Abcam 53087) were used for the study.

### Western Blot

Protein lysates from cultured cells were prepared with RIPA buffer containing protease inhibitors. 30 micrograms of lysate for each condition was separated on SDS/Polyacrylamide gels, transferred to PVDF membranes (Fisher Scientific, 88518) and incubated with antibodies following an earlier described protocol (29). Following antibodies were used for Western blots; p44/42 MAPK (9107, Cell Signaling Technology), Phospho (Tyr202/Tyr204) p44/42 MAPK (9101, Cell Signaling Technology), PKCζ (9372, Cell Signaling Technology), phospho (Thr410) PKCζ (32450, Cell Signaling Technology), c-MYC (9402, Cell Signaling Technology), β-Actin, (A5441, Sigma), CDC2 (77055, Cell Signaling Technology), Phospho (Tyr15) CDC2 (4539, Cell Signaling Technology). Histone H3 (39064, Active Motif), Phospho(Ser10)-Histone H3 (9701, Cell Signaling Technology). Phospho (Tyr416)-SRC Family (2101, Cell Signaling Technology). For secondary antibodies, Goat anti-mouse IgG-HRP (Santa Cruz, sc2005) and Goat anti-rabbit IgG-HRP ((Santa Cruz, sc2004) were used. Protein was visualized using chemiluminescence (ECL) substrate (Sigma, 689 WBLURO500) and imaged with a ChemiDoc system.

### Statistics and Reproducibility

To characterize the proteomes of hTSCs, STBs, and EVTs, we performed four independent biological replicates for each cell type. For all immunofluorescence and western blot, cell proliferation assays and RT-qPCR experiments, we performed at least three independent experiments. To generate RNA-seq data, we performed three independent experimental replicates in hTSCs, STBs and EVTs. The scRNA-seq data were generated from single cells isolated from two first-trimester placentas.

Statistical significances were calculated for hTSC proliferation analyses and RT-qPCR analyses. Independent datasets were analyzed using GraphPad Prism software. To determine statistical significance between control and experimental groups, we performed two-tailed, unpaired t test with Welch’s correction and indicated as mean ± SEM. Significantly altered values are highlighted (****p<0.0001, ***p<0.001, **p<0.01, *p<0.05).

## Data Availability

The raw proteomics data is submitted to MassIVE database (MSV000097626). Raw data for bulk RNA-seq in CT27 hTSCs are submitted to GEO database with accession numbers GSE296084 for hTSC stem state and EVT conditions and GSE214634 for STB condition. Raw data for scRNA-Seq in human first-trimester placentas are submitted with GEO accession number GSE145036.

## Results

### Identifying uniquely expressed proteins in human trophoblast cell types

We used Orbitrap mass analyzer for high-resolution, accurate-mass proteomics analyses with CT27 hTSCs, when cultured in stem state or after differentiation to STBs and EVTs. We validated differentiated fates by analyzing specific markers; HLA-G for EVT and hCG-β for STB differentiation (Fig. 1A). Proteins were identified and quantified from the raw data using Proteome Discoverer 3.0 with FDR cutoff 0.05. We considered expression of a protein when detected in at least two replicates of respective cell types. The combined global proteome analyses in all three-trophoblast cell types identified 393059 peptide sequences, corresponding to 4911 proteins (Fig. 1B). Pearson’s correlation coefficient analyses of replicates confirmed reproducible measurements in each trophoblast cell type (Fig. 1C). We identified 4542 proteins in hTSCs, 4518 proteins in STBs and 4559 proteins in EVTs. The distribution of identified proteins in each sample of three trophoblast cell types is shown in Figure 1D and mentioned in supplementary Dataset S1. Our analyses identified 4246 common proteins in all three trophoblast cell types (supplementary Dataset S1). The quality of the mass-spectrometry data was validated from selective enrichment of peptides from proteins that are known to be selectively expressed in specific cell types, such as LRP2 in hTSCs, hCG-β in STBs and HLA-G in EVTs (Fig. 1E).

**Figure 1:**
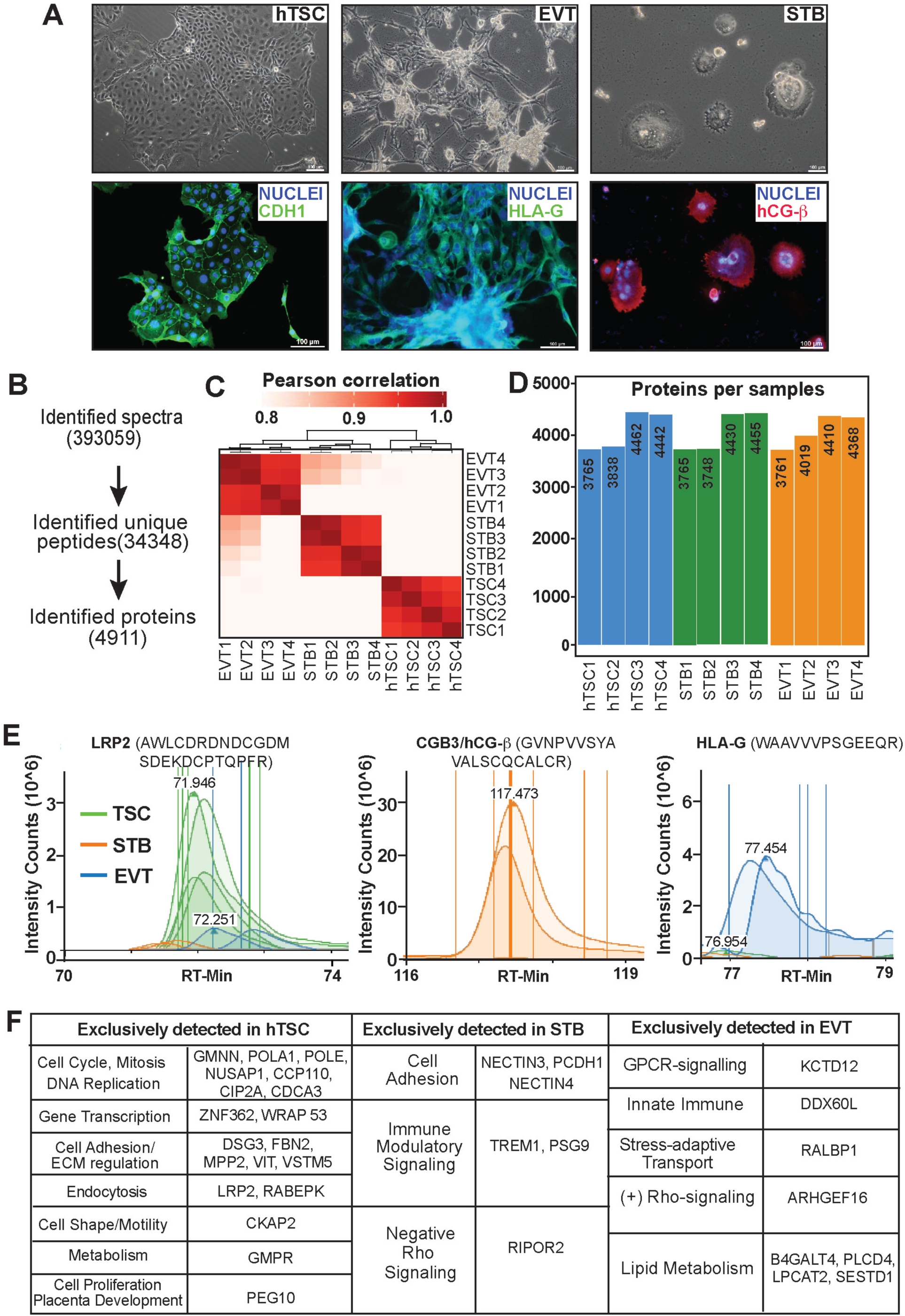
Global Proteomic Landscape in hTSCs and hTSC-derived STB and EVTs. (A) (Top panels), Cell images of hTSCs cultured in stem-state condition and after differentiation to STBs and EVTs. (Bottom panels) Immunofluorescence images showing CDH1, HLA-G and hCG-β expressions in TSC stem state, EVT and STBs, respectively, confirming specific cell fates. (B) Workflow for peptide and protein identifications. (C) Pearson’s correlation coefficient analyses of proteomes from experimental replicates of three trophoblast cell types. (D) Plot shows number of proteins identified in each replicate of three trophoblast cell types. (E) Examples of exclusively identified peptide spectrums in three trophoblast cell types. Indicated peptides from LRP2, hCG-b and HLA-G are exclusively identified in hTSCs, STBs and EVTs, respectively. (F) Exclusively identified proteins and their respective functions in hTSCs, STBs and EVTs are shown.

Next, we focused on proteins that are exclusively detected in hTSC stem state, in STBs and in EVTs. Initial screening identified 75 unique proteins in stem state, 77 unique proteins in STBs and 89 unique proteins upon EVT differentiation (supplementary Dataset S2). The undifferentiated hTSCs represent the CTB progenitors during early human placentation. Thus, to gain a better understanding of uniquely expressed proteins and their functions in three trophoblast cell types, we curated the proteomics data in hTSCs with single-cell RNA-seq data that we generated from first-trimester human placentae (Supplementary Fig. S1A, B) and reported earlier (28). Our curated analyses identified 19 proteins exclusively in undifferentiated hTSCs, 6 proteins exclusively in STBs and 8 proteins exclusively in EVTs (Fig. 1F). As expected, several of the 20 exclusively identified proteins in TSCs (GMNN, POLA1, POLE, NUSAP1, CCP110, CIP2A and CDCA3) are involved in DNA replication, cell cycle and mitosis. However, interestingly, the second major group of exclusively identified proteins in hTSCs is cell adhesion & extracellular matrix (ECM) proteins (DSG3, FBN2, MPP2, VIT and VSTM5). In addition, we also exclusively identified proteins in TSCs, which are implicated in gene regulation (ZNF362, WRAP53), endocytosis (LRP2, RABEPK), cell motility (CKAP2), and metabolism (GMPR). PEG10, a protein known to regulate placenta development (30) was also exclusively detected in TSCs (Fig. 1F).

NECTIN3, NECTIN4 and PCDH1, which were exclusively detected in STBs, are known to regulate cell-adhesion. Interestingly, NECTIN4, which mediates important functions at tight junctions, has been implicated as a predictive marker of Preeclampsia (31). We also identified STB-exclusive expression of RHO family interacting cell polarization regulator 2 (RIPOR2), a negative regulator of Rho-GTPase signaling pathway, and immune modulatory protein TREM1 and PSG9. In contrast, in EVTs, we exclusively determined expressions of ARHGEF16, a guanine nucleotide exchange factor for Rho family GTPases. Other proteins that were exclusively determined in EVTs are implicated in lipid metabolism (LPCAT2, PLCD4, B4GALT4, SESTD1), innate immune response (DDX60L), stress-adaptive transport (RALBP1) and G-protein–coupled receptor signaling (KCTD12).

### Integration of global gene and protein expressions to identify differentially expressed proteins during hTSC differentiation

Global proteomics analyses identified only limited number of proteins exclusively in hTSCs in stem-state and upon differentiation to STBs and EVTs. However, a large number of proteins that are expressed in hTSC in stem state are differentially expressed in STBs and EVTs. Thus, to further understand the dynamics of protein expression during STB and EVT development and to define the correlation between gene and protein expression patterns during human trophoblast development, we performed integrated analyses of global RNA expression (RNA-seq) with global protein expression. To that end, we first performed RNA-seq analyses in hTSCs in stem-state and after differentiation to STBs and EVTs. We identified differentially expressed genes during STB and EVT differentiation using DESeq2. Next, we wrote a Python script to integrate differentially expressed RNAs with differentially expressed proteins.

Setting a RNA expression change cut off at 1.5 fold, we identified 835 and 712 genes that were upregulated both at RNA and protein level during STB and EVT differentiation, respectively (Fig. 2A). In contrast, 1044 and 889 genes were downregulated both at RNA and protein levels during STBs and EVT differentiation, respectively (Fig. 2A, supplementary Dataset S3). We confirmed strong upregulation of protein expressions of many well-known STB and EVT markers, such as, CGA, PSG9, ENDOU (STB markers) and HLA-G, ERBB3, TEAD3 (EVT markers), upon hTSC differentiation (Fig. 2B, C). Whereas proteins specifically involved in cell cycle regulation such as CDK1, MKI67 and hTSC self-renewal process, such as, YAP1, were strongly downregulated. Not surprisingly, the gene set enrichment analyses (GSEA) showed that the protein expression of genes associated with cell cycle, mitosis, chromosome segregation were downregulated during both STB and EVT development (Fig. 2D, E). We also confirmed that proteins associated with cell-cell adhesion were induced during STB development. However, interestingly, the GSEA analyses indicated that both STB and EVT development is associated with induction of proteins that are implicated in immune response and defense responses to viruses.

**Figure 2:**
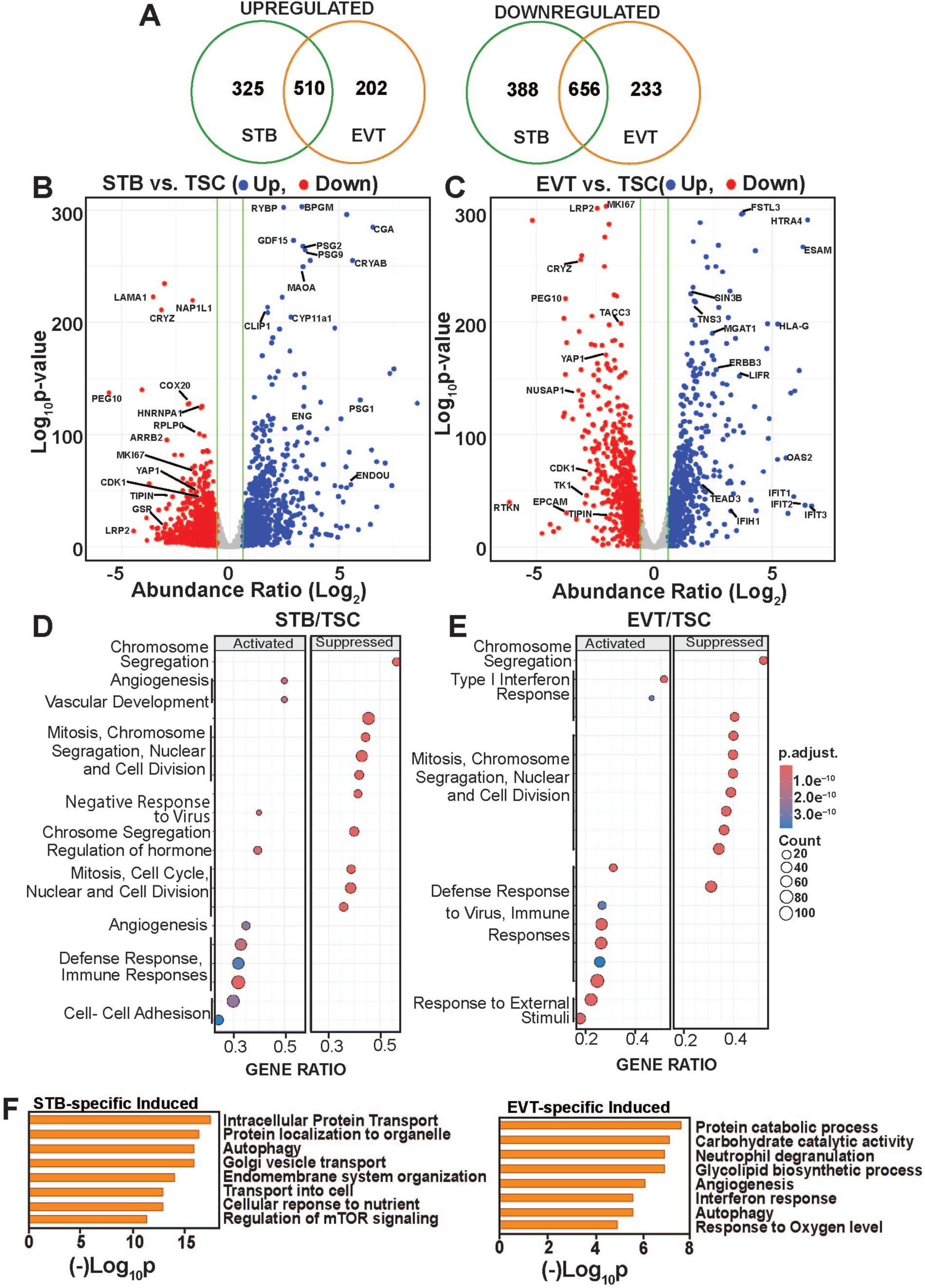
Global Protein Expression Dynamics During STB and EVT development. (A) Venn diagram showing number of differentially expressed proteins, identified via integrated RNA-seq and proteomic analyses, during STB and EVT differentiation. (B) and (C) Volcano plots of differentially expressed proteins during STB and EVT differentiation, respectively. Representative proteins with important functional implications are marked. (D) and (E) GSEA analyses of cellular functions of differentially expressed proteins during STB and EVT differentiation, respectively. (F) GO analyses of exclusively induced proteins during STB and EVT differentiation.

We identified several hundred proteins that were exclusively induced or repressed during STB and EVT differentiation (Fig. 2A and Supplementary Dataset S4). Gene Ontology (GO) analyses indicated that STB development were specifically associated with induction of proteins that are associated with cellular transport (ATP2B4, FOLR1, SLC7A1, SLC7A5, SLC12A4, SLC27A2, SLC38A2, SLC39A11, STRA6) and nutrient sensing mechanisms such as mTOR signaling, an important mechanism for fetal developmental programming (Fig. 2F). In contrast, EVT development is associated with upregulation of proteins that are associated with both ubiquitin–proteasome–mediated (TRIM21, UBE2L6, HERC4, RNF213), and lysosomal/autophagic protein catabolism (CTSL, TPP2, MAN1B1, DNAJC10, CHMP1B), carbohydrate metabolism (ALDOC, GAA, PCK2, PFKFB4), interferon responses (OAS1, IFI16, IFITM2), neutrophil degranulation (ADAM8, ADGRE5, COPB1, TSPAN14) and cellular responses to oxygen level (FOSL2, PLOD1) (Fig. 2F).

Autophagy is increasingly recognized as a key regulator of human trophoblast development. Physiological hypoxia induces autophagy in extravillous trophoblasts (EVTs), and inhibition of autophagy impairs EVT migration and invasion (32, 33). Studies using primary human trophoblasts have shown that autophagic flux is dynamically upregulated during syncytiotrophoblast differentiation and is required for efficient trophoblast fusion (34, 35). We noticed that both STB and EVT differentiations are associated with cell-type specific induction of proteins that are implicated in different stages of autophagy (Fig. 2F), such as autophagy initiation, autophagosome-lysosome fusion and autophagic degradation. In STBs, we identified upregulation of several ATG13, ATG4B, ATG3, WIPI2, WDR45, VPS18, CHMP6, CHMP4B, LAMP1 and CTSB, whereas in EVTs, we noticed induction of ATG9A, SH3GLB1, VPS39, CHMP1B, ATP6V1B1, VMA22 and CTSL. Our findings along with other studies indicate that autophagy could be a fundamental regulator of trophoblast development, coordinating both EVT differentiation and invasion as well as villous trophoblast syncytialization, thereby ensuring normal placental morphogenesis and function.

### Defining phosphoprotein dynamics during human trophoblast differentiation

One of the most important post-translational modifications regulating protein function is protein phosphorylation. However, phosphoproteomes in human trophoblast cells are largely unknown. Therefore, we analyzed the phosphoproteome in different trophoblast subtypes. We identified a total of 12355 phosphopeptides in our experimental samples. Among the 12355 phosphopeptides, more than 80% were singly phosphorylated, whereas 15.8% peptides were doubly phosphorylated and 3.7% peptides had more than 2 phosphorylation sites (Fig. 3A). Among the identified phosphosites, 91.2% were phosphorylated serine (S), 8.5% were phosphorylated threonine (T) and 0.3% were phosphorylated tyrosine (Y) (Fig. 3B). The reproducibility of our data set was demonstrated by pairwise Pearson correlation, which revealed close clustering of experimental replicates of each trophoblast cell type (Fig. 3C). To maximize the quality of our data set, we filtered out phosphosites that were detected at least two experimental replicates of each cell type. We identified a total of 2251 phosphoproteins in all three trophoblast cell types (supplementary Dataset S4). The identified phosphoproteins in each experimental samples are indicated in Figure 3D. Among 2251 identified phosphoproteins, 1709 were common in all three trophoblast cell types (Fig. 3E). We also exclusively identified 165 phosphoproteins in hTSC stem state, whereas, 21 and 49 phosphoproteins were exclusively identified in STBs and EVTs, respectively (Fig. 3E, supplementary Dataset S4).

**Figure 3:**
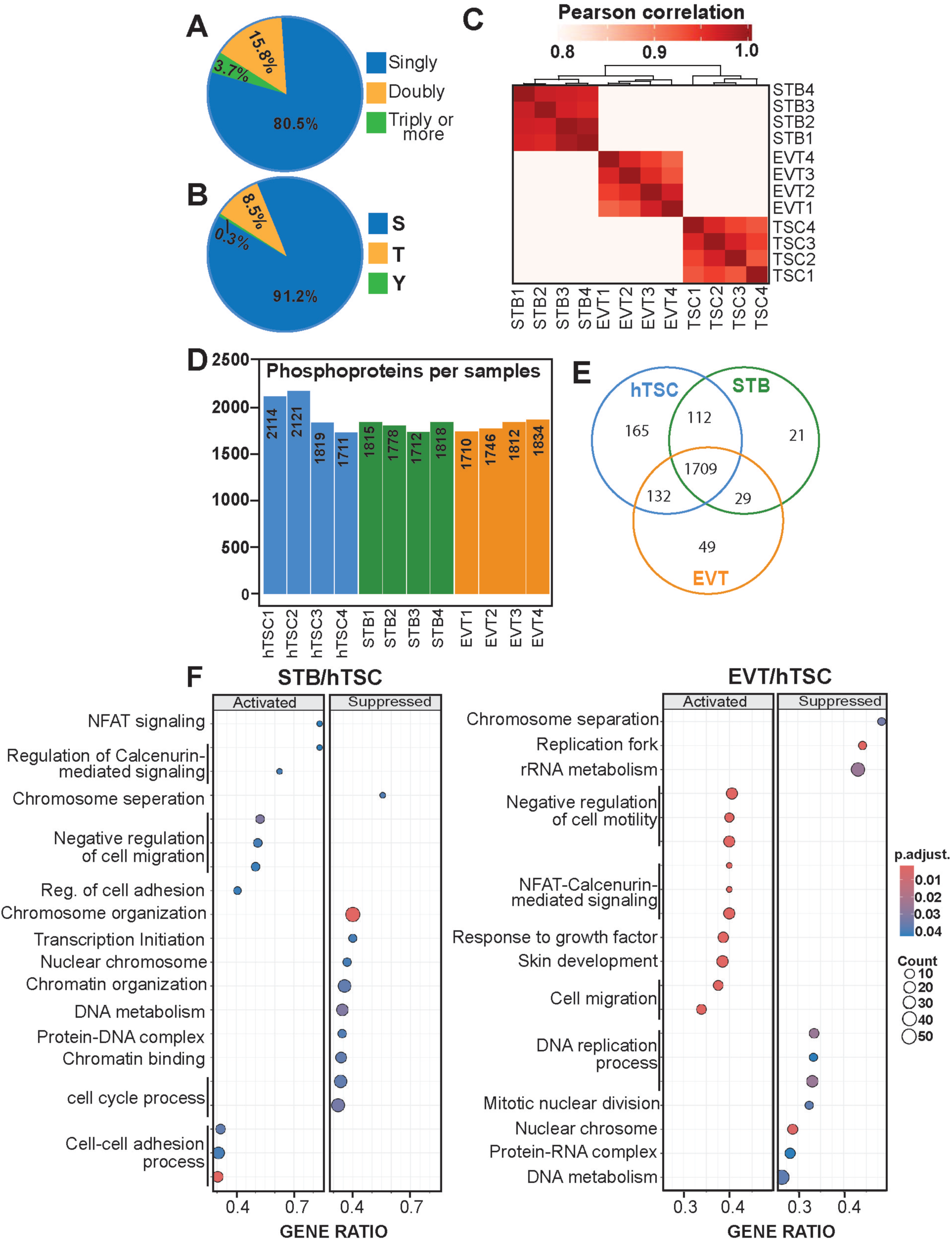
Global Phosphoprotein Dynamics in Human Trophoblast Cells. (A) Venn diagram showing percentage of all identified phosphopetides in three trophoblast cell types (hTSC, STB and EVT) with single, double and three or more phosphorylation events. (B) Venn diagram showing percentage of all identified phosphopeptides, which are phosphorylated on serine, threonine and tyrosine residues. (C) Pearson’s correlation coefficient analyses of phosphoproteomes from experimental replicates of three trophoblast cell types. (D) Plot shows number of phosphoproteins identified in each replicate of three trophoblast cell types. (E) Venn diagram showing number of uniquely identified as well as commonly identified phosphoproteins in hTSCs, STBs and EVTs. (F) GSEA analyses of cellular functions of differentially expressed phosphoproteins during STB and EVT differentiations.

Next, we performed pairwise analyses to identify differentially accumulating phosphosites during STB and EVT development. We identified 54 differentially accumulating phosphosites during hTSC to STB differentiation, of which 20 were upregulated in STBs and 44 were upregulated in hTSCs (supplementary Dataset S4). We also identified that hTSC to EVT differentiation is associated with 81 differentially accumulating phosphosites, of which 61 were upregulated in TSCs and 20 were upregulated in EVTs (supplementary Dataset S4). We next performed GSEA of exclusively and differentially phosphorylated proteins. The GSEA showed that the differential phosphorylation during both STB and EVT development led to suppression of cell cycle and mitosis. We also noticed that differential phosphorylation during STB development activates processes associated with cell-cell adhesion (Fig. 3F), whereas differential phosphorylation during EVT development strongly activated processes associated with cell motility.

Recently genome-wide CRISPR screening (12, 13) have identified essential regulators that promote hTSC stem-state and STB /EVT differentiation. We found that many essential regulators for stem-state maintenance (such as ARID3A, GATA3, EGFR, FOSL1, JUND) are phosphorylated in hTSCs (supplementary Dataset S4). We also found that some STB promoting (DNMT1, PBRM1, RPTOR, GRHL1, TFEB) and EVT promoting proteins (SPEN, POU2F1, FZR1) are also phosphorylated (supplementary Dataset S4).

### Identifying active protein kinases during human trophoblast development

The phosphoproteome within a cell is regulated by protein kinases, which in response to cell intrinsic or extrinsic signals convert substrate proteins to phosphorylated forms, thereby modulating cell-signaling processes. Thus, protein kinases are key regulatory components for cell fate decision. However, we have a very poor understanding of protein kinases that are essential for distinct stages of trophoblast development during human placentation. Therefore, we analyzed the phosphoproteome to identify specific protein kinase and their function in controlling hTSC stem-state and STB/EVT differentiation.

To identify active kinases in trophoblast cells, we used the human kinome from the KinHub database (36) and analyzed phosphoproteome data using RokaiXplorer computational tool. Next, we validated expressions of active kinases in respective trophoblast cell types of a developing human placenta using the scRNA-seq data from first-trimester human placenta. Curated list of active kinases in specific cell types is mentioned in supplemental dataset S5. As expected, the most active kinases identified in undifferentiated hTSCs are involved in cell cycle progression and cell proliferation, such as CDK1, CDK5, CDK6, and AURKA (Fig. 4A, B). We also identified MAPK13, a member of the p38 MAP kinase implicated in stress and inflammatory signaling pathway, is highly active in undifferentiated hTSCs. (Fig. 4A. B).

**Figure 4:**
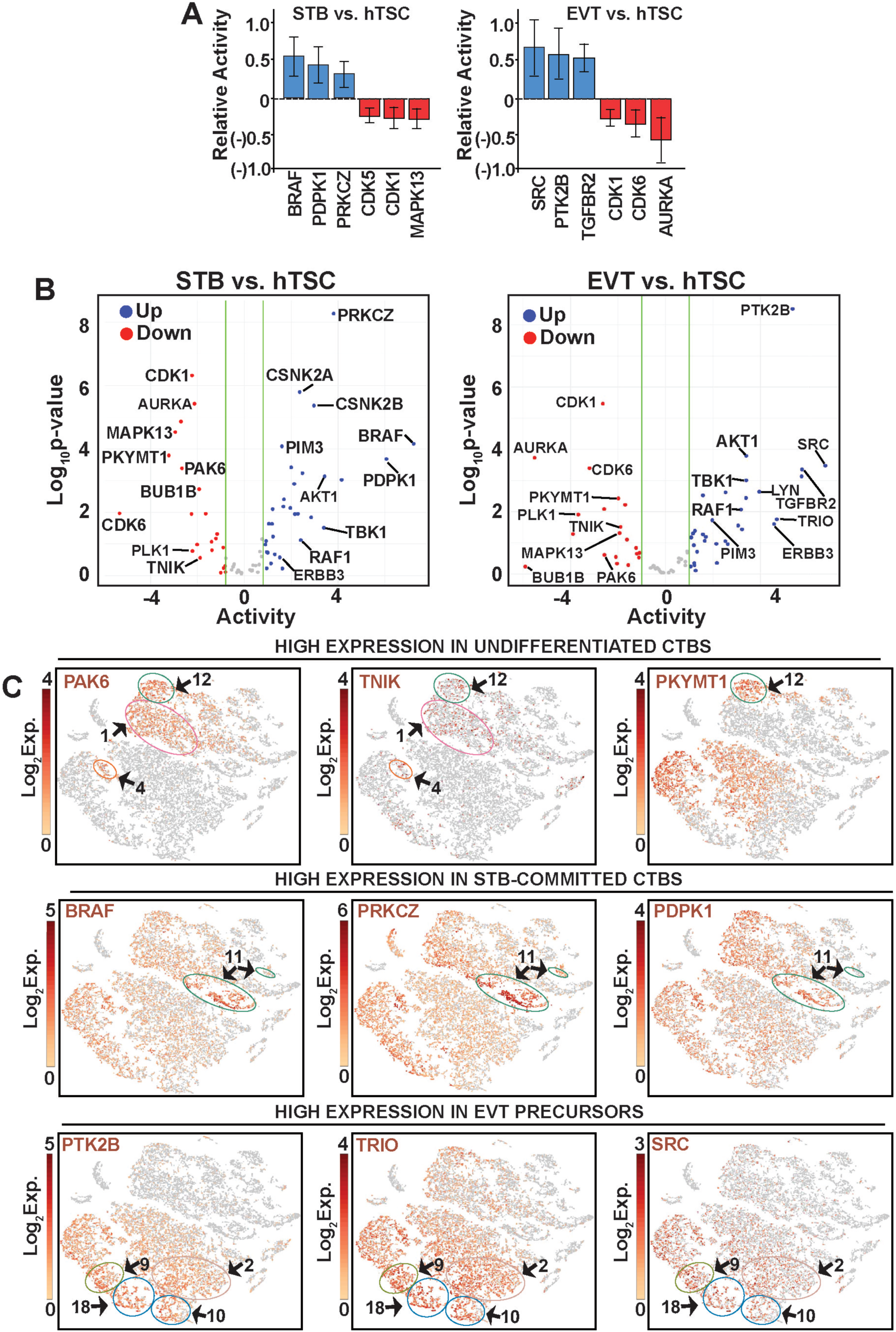
Identification of Active Kinases in Human Trophoblast Cells. (A) Plots show most differentially active kinases identified during STB and EVT differentiation. (B) Volcano plots of differentially active kinases during STB and EVT differentiation. Kinases implicated in different cell type-specific functions are indicated. (C) t-SNE Plots of scRNA-seq analyses in first-trimester placenta highlighting trophoblast cell-type specific mRNA expression patterns of different kinases.

Our analyses also identified many kinases that are selectively active in undifferentiated hTSCs. For example, membrane associated protein tyrosine/threonine kinase 1 (PKMYT1), which negatively regulates the G2/M transition (37) and BUB1 mitotic checkpoint kinase B (BUB1B), a serine/threonine kinase that delay the onset of anaphase to ensure proper chromosome segregation (38), are highly active in undifferentiated hTSCs (Fig. 4B, supplementary Dataset S5). Similarly, PAK6, a p21-stimulated serine/threonine protein kinase implicated in cytoskeleton rearrangement (39), and TRAF2 and NCK interacting kinase (TNIK), another serine/threonine kinase and an activator of the WNT signaling pathway (40), are also highly active in hTSCs. The scRNA-seq analyses data also confirmed that mRNA expressions of all of these kinases are highly enriched in undifferentiated CTBs of a developing first-trimester human placenta (Fig. 4C).

We also identified kinases that are selectively active in STBs and EVTs. We found that B-Raf proto-oncogene (BRAF), 3-phosphoinositide dependent protein kinase 1 (PDPK1) and atypical protein kinase C isoform, protein kinase zeta (aPKCζ, encoded by the gene *PRKCZ*) are highly active kinases in STBs (Fig. 4A, B, supplementary Dataset S5). The scRNA-seq analyses confirmed that expressions of all of these kinases are induced in CTBs, which are committed to STB development (Fig. 4C). High activity of casein kinase 2 subunits (CSNK2A1, CSNK2A2 and CSNK2B), which regulate various cellular processes including metabolism, was also detected in STBs (supplementary Dataset S5). Active kinases, selectively identified in EVTs, include tyrosine-Protein Kinase Src (SRC), Protein Tyrosine Kinase 2 Beta (PTK2B, also known as PYK2, a non-receptor tyrosine kinase, implicated in focal adhesion) and Transforming Growth Factor Beta Receptor 2 (TGFBR2) (Fig. 4A, 4B). Another SRC family tyrosine kinase, LYN, which is implicated in CTB syncytialization in human (41) and trophoblast giant cell development in rodents (42), and TRIO, a Rho guanine nucleotide exchange factor, implicated in cell migration (43) are also highly active in EVTs. These kinases are also highly expressed in EVT precursors of a developing human placenta (Fig. 4C). In addition to these cell-specific kinase activity, we also identified many kinases, such as AKT1, ERBB2, ERBB3, RAF1 and TBK1, which are active in both STBs and EVTs (Fig. 4B, and supplementary Dataset S5).

### Testing functional importance of kinases in hTSCs

Many of the kinases that are highly expressed and active in undifferentiated CTB/hTSCs, such as PKMYT1, BUB1B, PAK6, and TNIK are implicated in various cancers and are targeted for cancer treatment (44–49). Similarly, kinases that are induced and active in STBs (such as BRAF and aPKCζ) and EVTs (such as SRC, PTK2B and TRIO) are implicated in various cancers and are therapeutic targets (50–54). However, it is unknown whether development and function of human trophoblast cells are affected upon inhibition of these kinases. Therefore, we tested whether maintenance of hTSC stem-state or their differentiation to STBs and EVTs are affected upon pharmacological inhibition of these kinases. Using selective pharmacological inhibitors (Table 1), we tested importance of PAK6, TNIK, BUB1B and PKMYT1 in undifferentiated hTSCs, importance of aPKCζ in STB development and importance of SRC, PTK2B, TRIO and LYN in EVT development.

We used GNE2861 (a selective inhibitor of group II p21-activated kinases, including PAK6), Rentosertib (TNIK inhibitor), Antitumor agent 180 (ATA180, BUB1B inhibitor), Lunresertib (PKMYT1 inhibitor), Go6983 (aPKCζ inhibitor), SKI1 (SRC inhibitor), Bafetinib (LYN inhibitor), PF4618433 (PYK2 inhibitor) and ITX3 (TRIO inhibitor) for our study. Because inhibitor activities had not been established in hTSCs, we first optimized working concentrations using MTT assays and selected doses (Table 1) that maintained >80% hTSC viability, except for SKI1 and PF4618433, which retained ∼72% and ∼77% viability, respectively, when tested with Annexin-V-FITC apoptosis assay in stem-state culture condition (Supplementary Fig. S3).

We validated inhibition of PAK6, TNIK, BUB1B and PKMYT1 activity in presence of respective inhibitors by testing their downstream effectors. PAK family kinases including PAK6 promote p42-44 MAPK (ERK1/2) phosphorylation (55, 56). We validated that ERK1/2 phosphorylation is inhibited in hTSCs when treated with GNE2861 (Fig. 5A). TNIK is a regulatory component of the β-catenin/T-cell factor 4 transcriptional complex, and its function is important to induce Wnt/ β-catenin target genes, such as c*MYC* (49, 57). Thus, we validated TNIK inhibition by testing loss of cMYC expression upon treating hTSCs with Rentosertib (Fig. 5A). To validate inhibition of BUB1B function, we tested Histone H3 phophorylation at serine 10 (pH3S10), a marker for mitosis. Through its role at kinetochores, BUB1B helps to regulate the chromosome passenger complex and thereby influences the activity and localization of Aurora kinase B, the principal kinase responsible for phosphorylating pH3S10 modification (58–60). Accordingly, pharmacological inhibition of BUB1B is expected to disrupt Aurora B-dependent mitotic signaling indirectly. Consistent with this, ATA180 treatment reduces H3S10 phosphorylation in hTSCs (Fig. 5A). PKMYT1 regulates cell cycle by phosphorylating CDC2 on both threonine-14 and tyrosine-15 (61). We confirmed loss of phspho-tyrosine15 CDC2 (pY15CDC2) when hTSCs were treated with Lunresertib (Fig. 5A).

**Figure 5:**
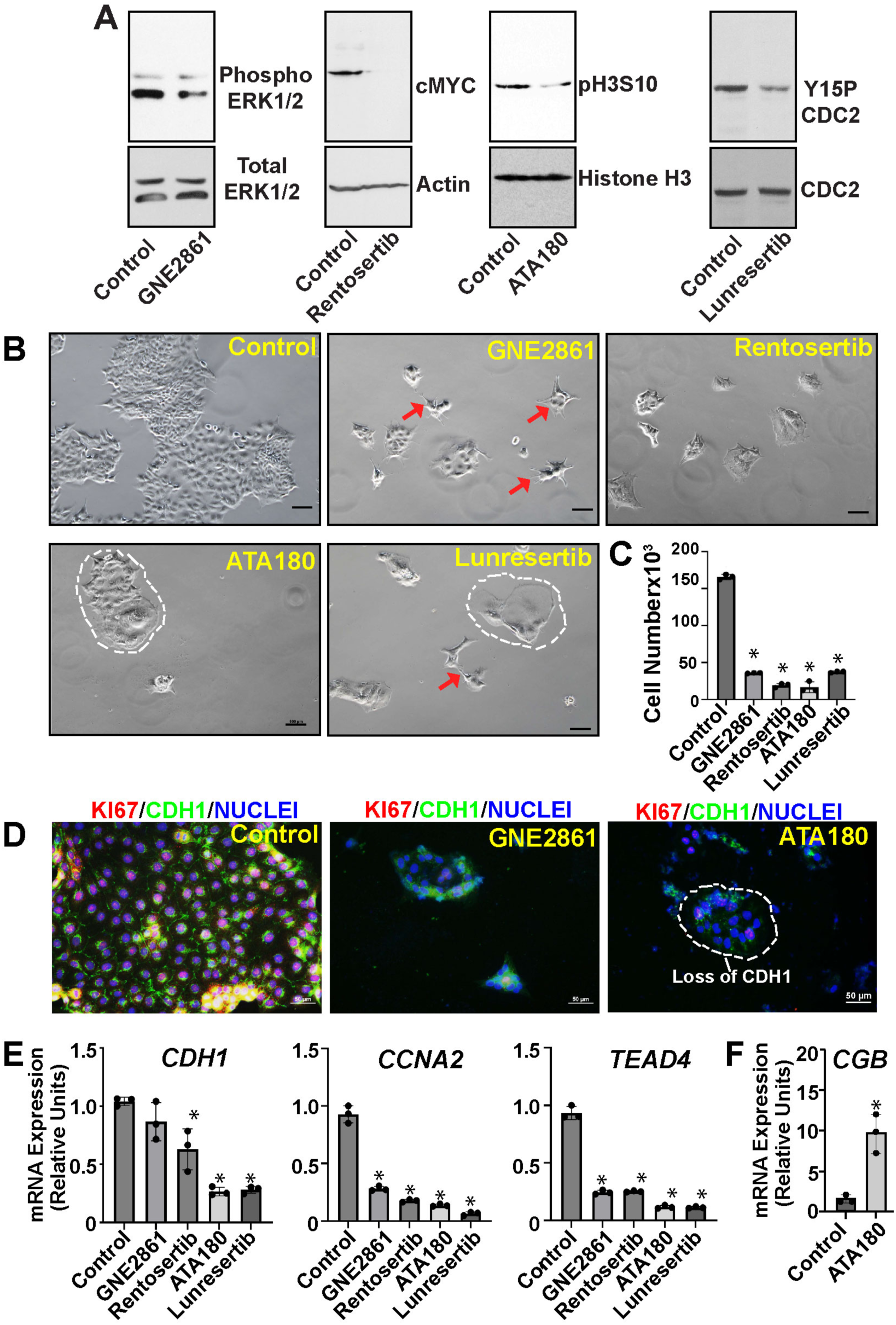
Pharmacological Inhibition of PKMYT1, BUB1B, PAK6 and TNIK affects hTSC proliferation and expression of stem-state genes. (A) Inhibition of hTSC stem-state kinase activities were validated by monitoring established downstream signaling events. Representative western blots are shown. DMSO (vehicle)-treated hTSCs were used as control. GNE2861 reduced ERK1/2 phosphorylation, consistent with inhibition of PAK6-dependent MAPK signaling. Rentosertib suppressed cMYC expression, confirming inhibition of TNIK-mediated Wnt/β-catenin signaling. ATA180 decreased pH3S10 level, reflecting disruption of BUB1B-dependent mitotic signaling through Aurora B kinase, whereas Lunresertib abolished inhibitory CDC2 Tyr15 phosphorylation, consistent with PKMYT1 inhibition. (B) Micrographs show hTSC colony morphologies when cultured in presence of different kinase inhibitors. DMSO-treated hTSCs were used as control. Red arrows indicate that hTSCs adapted an invading phenotype with visible cell protrusions, a morphological change often noticeable when hTSCs undergo EVT differentiation process. White borders indicate hTSC colonies with flattened morphology and apparent losses of individual cellular boundaries indicating adaptation to STB-like fate. (C) The plot shows significant reduction in hTSC proliferations upon culturing with inhibitors for 72h. Data are representative of 4 independent experiments (n=4). Statistical significances were calculated for each inhibitor condition with respect to the control and indicated as mean ± SEM. Two-tailed unpaired t-test with Welch’s correction, *P≤0.01. (D) Immunofluorescence imaging showing reduced KI67 expression in TSC colonies, treated with GNE 2861 (PAK6 inhibitor) and ATA180 (BUB1B inhibitor). Note loss of CDH1 (cellular boundaries) in ATA180 treated cells. (E) RT-qPCR analyses showing loss of mRNA expressions of hTSC stem-state genes upon treatment with kinase inhibitors. DMSO-treated cells were used as control. Gene expression levels are plotted as relative units with respect to expression levels in untreated CT27 hTSCs. Data are representative of 3 independent experiments (n=3). Statistical significances were calculated for each inhibitor condition with respect to the control and indicated as mean ± SEM. Two-tailed unpaired t-test with Welch’s correction, *P≤0.01. (F) RT-qPCR analyses showing induction of STB-specific gene *CGB,* when hTSCs were treated with ATA180 in stem-state culture condition. Gene expression levels are plotted as relative units with respect to expression levels in untreated CT27 hTSCs. Data are representative of 3 independent experiments and indicated as mean ± SEM. Two-tailed unpaired t-test with Welch’s correction, *P≤0.01.

Inhibition of PAK6, TNIK, BUB1B, or PKMYT1 markedly impaired hTSC proliferation, as demonstrated by reduced cell growth and Ki67 expression (Fig. 5B–D). Expression of stem-state regulators, including *CDH1, CCNA2*, and *TEAD4*, was significantly reduced following inhibition of each kinase (Fig. 5E), indicating loss of the hTSC stem/progenitor state. Distinct morphological changes accompanied these molecular defects. PAK6 inhibition induced invasive cellular protrusions resembling early EVT differentiation (Fig. 5B, red arrows). BUB1B inhibition produced flattened colonies with loss of cell-cell boundaries and reduced CDH1 expression, accompanied by induction of the STB marker *CGB*, indicating adaptation to STB-like fate (Fig. 5B,D,F). Inhibition of TNIK or PKMYT1 resulted in mixed invasive and flattened morphologies, consistent with aberrant lineage commitment. Together, these findings identify PAK6, TNIK, BUB1B, and PKMYT1 as essential regulators of hTSC proliferation and maintenance of the trophoblast stem/progenitor state.

When maintained in stem-state culture condition, hTSC proliferation and stem-state morphology were not affected (Supplementary Figure S4A) in the presence of Go6983, which inhibits atypical PKC isoforms aPKCζ and aPKCλ/i (52, 62). RT-qPCR analyses showed that mRNA expressions of hTSC stem-state marker, *TEAD4* and STB marker *CGB* were not significantly altered (Supplementary Figure S4B). In contrast, STB differentiation was severely affected in Go6983-treated hTSCs. Under STB differentiation condition, Go6983-treated hTSCs maintained a stem-state like morphology (Fig. 6A). Western blot analyses confirmed that aPKCζ phosphorylation at Threonine 410 (T410P aPKCζ) was inhibited upon Go6983 treatment (Fig. 6B). The Go6983-treated cells maintained high-level mRNA expressions of stem state regulators, such as *CDH1, TP63,* and *TEAD4*, under STB differentiating condition, whereas, induction of STB markers, such as *ERVW1, SDC1, and CGB* were significantly inhibited (Fig. 6C, D). These findings indicate aPKCζ activity as a critical regulator of human STB differentiation while demonstrating that it is dispensable for hTSC stem-state maintenance. A recent report (63), which showed that aPKCζ-III, an isoform of aPKCζ, promotes fusion of CTB progenitors to form STBs, further support this aspect.

**Figure 6:**
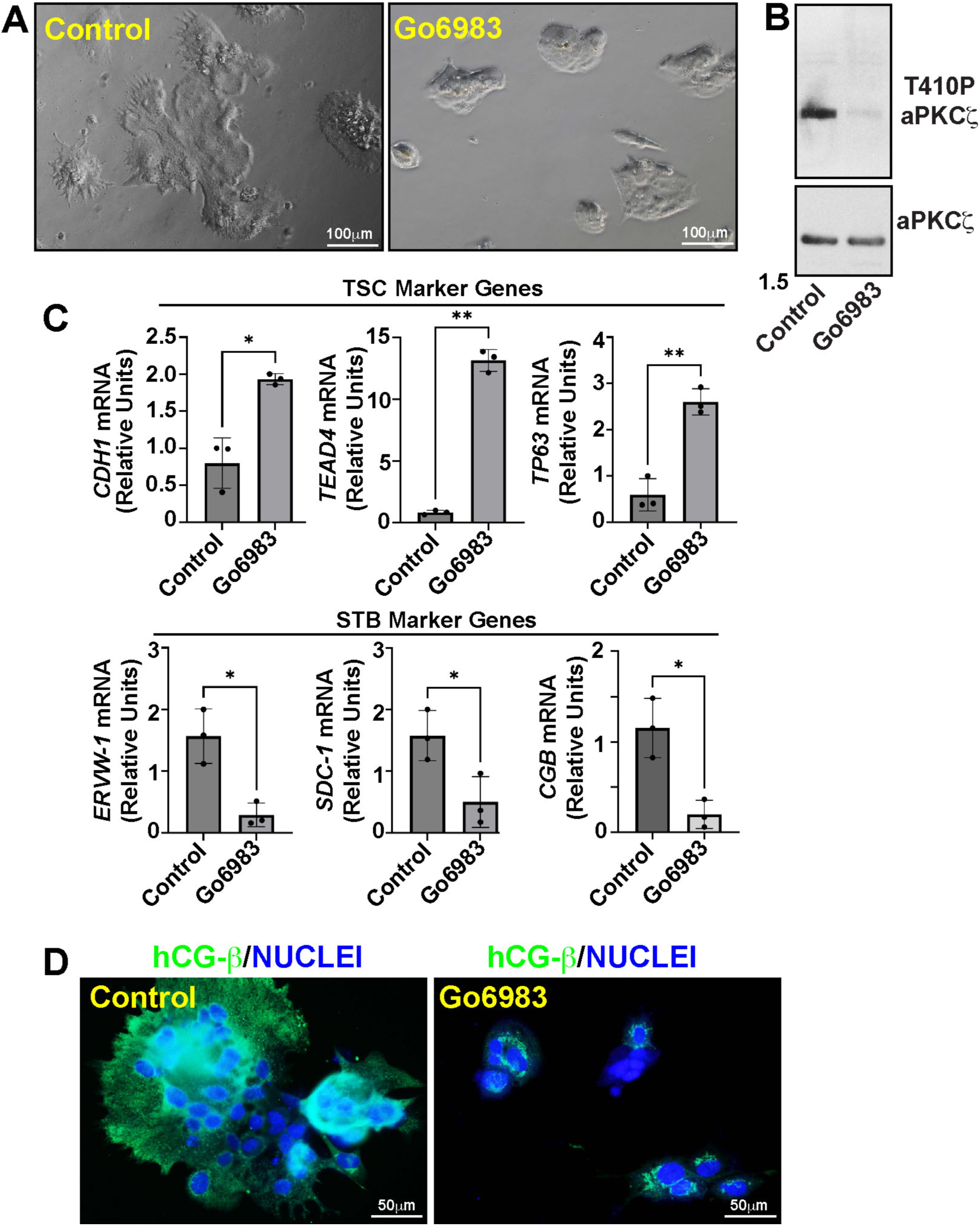
Pharmacological Inhibition of aPKC isoforms prevents STB differentiation in hTSCs. (A) Micrographs show hTSCs after culturing with or without Go6983 for 5 days in a culture condition that promotes STB differentiation. DMSO (vehicle)-treated hTSCs were used as control. (B) Representative Western blots confirming inhibition of aPKCζ activation by loss of Thr410 phosphorylation. (C) RT-qPCR analyses showing high-level mRNA expressions of hTSC stem state genes and low-level mRNA expressions of STB-specific genes in Go6983-treated hTSCs compared to DMSO-treated hTSCs (Control), when cultured in STB culture condition. Gene expression levels are plotted as relative units with respect to expression levels in untreated CT27 hTSCs after STB differentiation. Data are representative of 3 independent experiments and indicated as mean ± SEM. Two-tailed unpaired t-test with Welch’s correction, *P≤0.01. (D) Representative immunofluorescence images confirming reduced hCG-b protein expression in Go6983-treated cells.

Next, we tested whether kinases identified as active in EVTs are required for hTSC stem-state maintenance and EVT development. Under stem-state culture conditions, inhibition of TRIO with ITX3 or LYN with bafetinib did not significantly affect hTSC proliferation or stem-state maintenance, whereas inhibition of SRC with SKI1 and PTK2B with PF-4618433 partially reduced hTSC proliferation (Supplementary Fig. S4A). In contrast, EVT differentiation was strongly impaired by all four kinase inhibitors. Under EVT-inducing conditions, control hTSCs acquired characteristic EVT-like morphology with extensive cellular protrusions, whereas inhibitor-treated cells showed defective morphology and reduced protrusive outgrowth. Notably, bafetinib-treated cells appeared unhealthy and showed increased apoptosis (Fig. 7A; Supplementary Fig. S4C).

**Figure 7:**
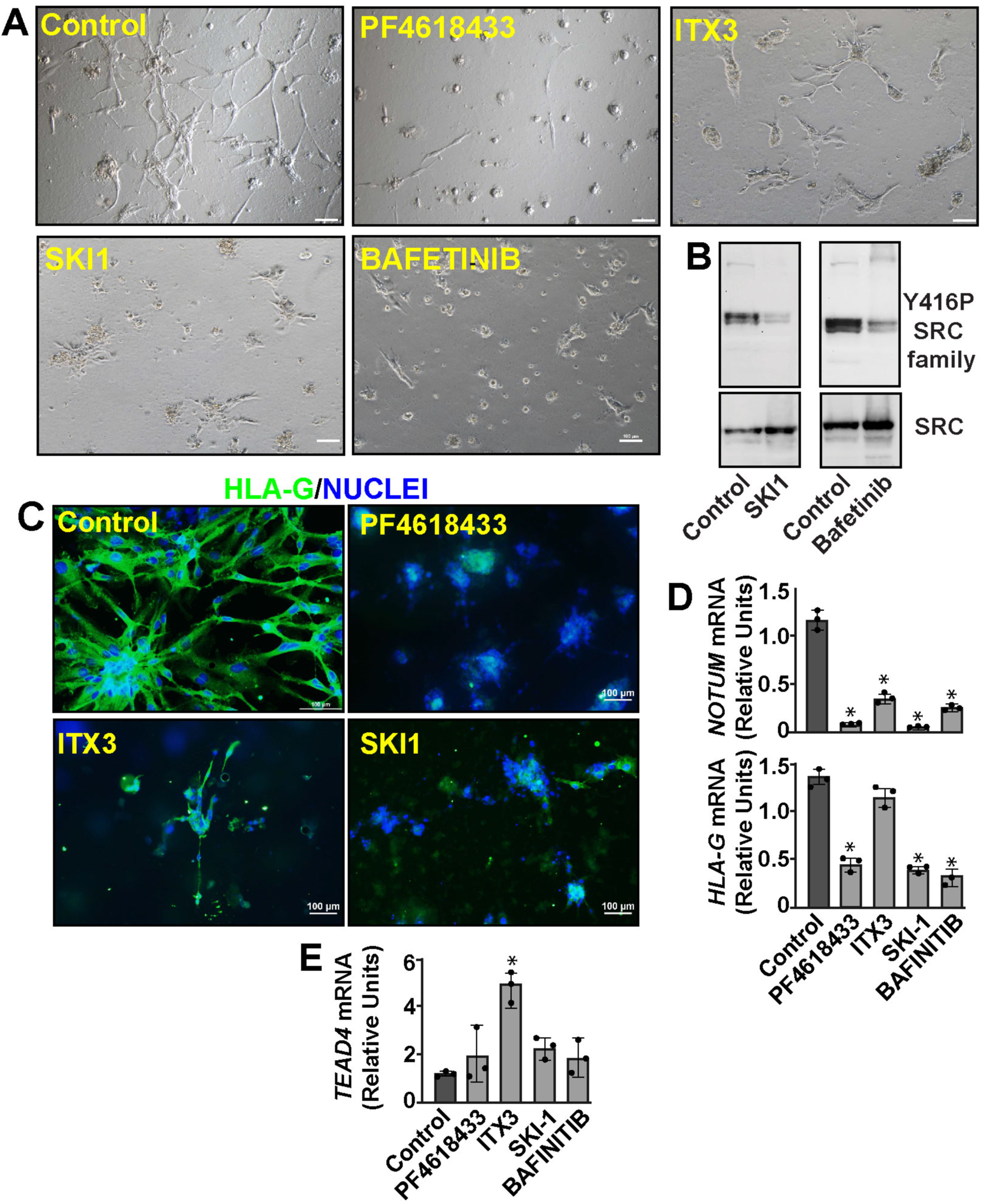
Pharmacological Inhibition of SRC, PTK2B, TRIO and LYN negatively affect EVT differentiation in hTSCs. (A) Micrographs show hTSCs after culturing with or without kinase inhibitors for 6 days in a culture condition that promotes EVT differentiation. DMSO (vehicle)-treated hTSCs were used as control. (B) Representative Western blots showing inhibition of either SRC or LYN reduced Y416PSRC-family signal, supporting suppression of SRC-family kinase activity under these conditions. (C) Representative Immunofluorescence images confirming reduced HLA-G protein expression in kinase inhibitor-treated cells. (D) RT-qPCR analyses showing reduced mRNA expressions of EVT-specific genes *NOTUM* and *HLA-G* in kinase inhibitor-treated cells compared to DMSO-treated hTSCs (Control), when cultured in EVT culture condition. Gene expression levels are plotted as relative units with respect to expression levels in untreated CT27 hTSCs after EVT differentiation. Data are representative of 3 independent experiments and indicated as mean ± SEM. Two-tailed unpaired t-test with Welch’s correction, *P≤0.01. (E) RT-qPCR analyses showing maintenance of high-level mRNA expression of *TEAD4*, a hTSC stem state gene, in cells cultured in EVT condition with ITX3, a selective TRIO kinase inhibitor. *TEAD4* expression was not significantly induced with other kinase inhibitors. Gene expression levels are plotted as relative units with respect to expression levels in untreated CT27 hTSCs after EVT differentiation. Data are representative of 3 independent experiments and indicated as mean ± SEM. Two-tailed unpaired t-test with Welch’s correction, *P≤0.01.

Because phosphorylation of SRC at Tyrosine 416 is essential for SRC kinase activation and occurs predominantly through trans-autophosphorylation (64–67), we assessed phospho-SRC family Tyr416 levels as a functional readout of SRC-family kinase activity. LYN, another SRC-family kinase, is similarly activated by autophosphorylation of its activation-loop tyrosine, Tyrosine 396/397, which is homologous to SRC Tyrosine 416 (68). Consistent with this conserved activation mechanism, inhibition of either SRC or LYN reduced phospho-SRC-family Tyrosine 416 (Y416P SRC Family) signal (Fig. 7B), supporting suppression of SRC-family kinase activity under these conditions. However, because suitable antibodies were not available, we could not directly confirm inhibition of TRIO or PTK2B at the protein-phosphorylation level.

We further validated defective EVT development by assessing EVT markers, including HLA-G and NOTUM, which were robustly induced under control EVT-differentiation conditions but were inefficiently induced following kinase inhibition (Fig. 7C,D). We also examined expression of the stem-state gene *TEAD4* and observed significant *TEAD4* upregulation only upon TRIO inhibition (Fig. 7E). Together, these findings indicate that SRC, PTK2B, TRIO, and LYN activities are required for efficient EVT development during human placentation, whereas SRC and PTK2B also contribute to maintenance of the hTSC/CTB stem-progenitor state.

## Discussions

hTSC cell culture model largely recapitulates the gene expression programs of the primary trophoblast cells of a developing human placenta (11, 26, 28, 69). Thus, this is a valuable model to study mechanisms that guide trophoblast cell lineage development and function during human placentation. In this study, we used an integrated proteomics and genomics approach to definitively identify protein expression dynamics during human trophoblast development. The integration of our phosphoproteomics data along with scRNA-seq data from primary trophoblast cells of a developing human placenta provided a stringent platform to identify active kinases that dictate trophoblast cell-specific functions during human placentation. Finally, experimental validation of importance of specific kinases in hTSC stem-state maintenance as well as in STB/EVT differentiation solidified our findings from the integrated omics approach. Thus, our findings in this study will be an important resource for future understanding of kinase-mediated processes in human trophoblast cells.

Cellular phosphoproteome is regulated by the coordinated activities of both kinases and phosphatases. Consistent with this, our proteomic analyses identified several phosphatases that are differentially expressed during human trophoblast development. For example, PTPN12 is upregulated, whereas PTPN1 is downregulated during both STB and EVT differentiation, while PTPN23 is selectively enriched in EVTs (Supplementary Dataset S3). Although these findings suggest that phosphatases may also play important roles in regulating trophoblast lineage specification and function, the primary objective of the present study was to define the trophoblast kinome and phosphoproteome and to functionally characterize selected kinases. Accordingly, mechanistic investigation of individual phosphatases is beyond the scope of the current study and will be an important direction for future work.

The gene expression dynamics, high proliferation rate and self-renewal ability of undifferentiated hTSCs recapitulate the characteristics of proliferating CTB progenitors of developing human placenta (28). Thus, as expected, global proteome, phosphoproteome and kinase activity in undifferentiated hTSCs mostly indicated proteins and kinases that are involved in cell-cycle progression and mitosis. However, interestingly, we also exclusively identified a group of proteins in hTSCs, which are involved in cell-adhesion, such as DSG3 (Desmoglein 3) (70), extracellular matrix regulation, such as FBN2 (Fibrillin2), MPP2 (MAGUK P55 scaffold protein 2) and membrane proximal signaling, [vitrin (VIT) and V-Set and transmembrane domain containing 5 (VSTM5)]. Functional significances of these proteins have been implicated in other cellular contexts. FBN2 is implicated in macular degeneration (71), MPP2 in synaptic adhesion (72), VIT in cartilage development (73) and Brain asymmetry (74) and VSTM5 in neuronal morphology and dendrite formation (75) and Vogt–Koyanagi–Harada disease (76). However, their functional importance in hTSCs/CTBs is yet to be identified.

Active WNT signaling is important to maintain hTSC stem state. Thus, we posited that inhibition of TNIK, a kinase involved in WNT signaling pathway, would promote differentiation. The adaptation of both STB-like and EVT-like cellular morphology upon TNIK inhibition validated our assumptions. We also noticed extensive cell protrusion upon PAK6 inhibition in hTSCs, indicating adaptation of a migratory phenotype. Intriguingly, noticeable transition from hTSC to STB like colony morphology were also observed upon inhibition of PKMYT1 and BUB1B, which are implicated in inhibitory regulation of CDK1 and proper chromosome segregation, respectively, during cell cycle process. During human placentation CTB progenitors adapt two distinct phenotypes with completely opposite cellular morphology, the syncytialized STBs and invasive EVTs. Collectively, our findings indicate that functions of these kinases in CTB progenitors might prevent premature adaptation of STB and EVT-fate, thereby balancing the trophoblast development leading to proper placentation.

Earlier, we reported that genetic depletion of atypical PKC isoform aPKCλ/i inhibits STB differentiation in hTSCs (69). Our proteomics study detected aPKCλ/i expression in all three trophoblast cell types. In contrast, aPKCζ expression is highly upregulated in STBs and impairment of STB development upon Go6983 indicates that aPKCζ function is important in STB development. We exclusively identified protein expression of RIPOR2 in STBs. RIPOR2 directly binds to RHO proteins to negatively regulate RHO-mediated cellular functions, including cell motility (77). RIPOR2 is also implicated in establishing and maintaining cell polarity. The aPKC isoforms are also important to maintain cell polarity. Maintenance of cell polarity in STBs is essential for nutrient transport from maternal blood to fetus (78, 79). Thus, along with aPKCλ/i and aPKCζ, RIPOR2 function could be crucial for STB function and embryonic development. Interestingly, PDPK1, which acts downstream to PI3 kinase (PI3K) (80) is highly active in STBs and is known to activate aPKCζ. Our scRNA-seq analyses showed that PDPK1 mRNA expressions is highly induced during CTB to STB transition and PI3K subunits are expressed in both undifferentiated and differentiating CTBs. Thus, we predict that a PI3K-PIP3-PDPK1-aPKCζ signaling axis is important for STB development. Our analyses also showed that several other kinases, including BRAF, FYN and EPHB4 are highly active in STBs and scRNA-seq analyses showed that their mRNA expressions are highly induced during CTB to STB transition. Thus, future studies to test their importance in STB development and function could provide important insight regarding human placentation.

One interesting aspect that we noticed during EVT development is induction of proteins associated with interferon/antiviral response, such as, OAS1, OAS2, IFIT2, IFIT3 and IFIH1. However, the mechanism that induces these interferon response proteins and their functional importance in EVTs are unknown. Human trophoblast cells harbor many endogenous retroviral elements, which have been implicated in trophoblast gene regulation (81, 82). Thus, it is possible that EVT development is associated with selectively enhanced expression of retroviral elements leading to induction of interferon response genes. However, other molecular causes such as degradation of mitochondrial DNA (83) or other cell-intrinsic signaling might activate the cyclic GMP–AMP synthase–stimulator of interferon genes (cGAS–STING) pathway to induce interferon response genes. To that end, the induction and activity of LYN kinase during EVT development is of further interest. In other cellular contexts, Lyn inhibits inflammasome formation by phosphorylating NLRP3 (84). Thus, the induction of interferon response genes and activity of LYN could be correlated during EVT development.

TGFβ signaling has been implicated in EVT development (85). Not surprisingly, TGFBR2 is one of the most abundant kinases in EVTs. However, we found that EVT development is also dependent on functions of SRC, PTK2B and TRIO. The tyrosine kinase SRC mediates multitude of cellular functions. In contrast PTK2B, a non-receptor tyrosine kinase, and TRIO, a Rho family guanine nucleotide exchange factor, are implicated in cell migration (43, 86). PTK2B is also known to regulate epithelial to mesenchymal transition (EMT) (87). The EVT development is associated with activation of several EMT regulators and adaptation of an invasive phenotype. Thus, defective EVT development upon PTK2B and TRIO inhibition could be due to defect in activity of EMT regulators along with mechanisms that promote cell invasion.

In addition to cell type specific kinases, we identified many kinases that are simultaneously active in multiple cell types. For example, we identified that several MAP kinase family members, such as MAPK14, MAP4K4, MAPK3 and MAPK8, are active in both undifferentiated hTSCs and in EVTs. The global RNA-seq, scRNA-seq and proteomics data confirmed that these kinases are highly expressed in CTBs/hTSC and EVTs. Likewise, AKT1, ERBB2 and ERBB3 are identified to be active in both STBs and EVTs. We also noticed that kinases with similar functions are selectively expressed within a specific trophoblast cells. For example, serum/glucocorticoid regulated kinase family member 3 (SGK3) is selectively induced in CTB progenitors and undifferentiated hTSCs. In contrast SGK1 is selectively induced and active in STBs. Notably, the receptor tyrosine kinase EPH receptor B4 (EPHB4) is highly active in STBs, whereas EPHB3 is highly active in EVTs. Thus, in future, it will be interesting to define how different kinases of same family members exert cell-type specific functions during human placentation.

Our findings have important translational implications for pregnancy-associated disorders characterized by defective trophoblast development, including recurrent pregnancy loss, preeclampsia, fetal growth restriction, and preterm birth. Many of the kinases identified in this study are already under active investigation as therapeutic targets in oncology, and highly selective small-molecule inhibitors have been developed for several of these pathways. Our data demonstrate, however, that these kinases also perform essential lineage-specific functions during normal human placentation, highlighting the need for caution when considering the use of kinase-targeted therapies during pregnancy. Conversely, defining the kinase signaling networks that govern trophoblast stem-state maintenance and lineage specification provides a framework for identifying dysregulated pathways in high-risk pregnancies and may facilitate the development of precision therapeutic strategies aimed at restoring, rather than suppressing, trophoblast signaling. As technologies for placenta-targeted drug delivery continue to advance, selective modulation of disease-associated kinase pathways may ultimately provide new opportunities to improve placental function while minimizing fetal exposure.

## Author Contributions

Soumen Paul conceived the project, designed experiments and wrote the manuscript. Rajnish Kumar performed all data processing and downstream bioinformatics analyses and wrote the initial version of the manuscript. Purbasa Dasgupta designed and performed experiments, analyzed data. Soma Ray worked with kinase inhibitors and hTSCs.

## Acknowledgement

This research was supported by NIH grants HD113673, HD103161, HD062546, HD101319, HD119510 and Next Gen Pregnancy Initiative Burroughs Wellcome Fund to S.P. This study was also supported by various core facilities, including the Mass spectrometry core, Genomics Core, Flowcytometry core, Imaging and Histology Core facility of the University of Kansas Medical Center. We acknowledge support from Asef Jawad Niloy and Mounika Vallakati for general experimental support. We also acknowledge administrative support from the Department of Pathology Internal Medicine and the Institute for Reproduction and Developmental Sciences of the University of Kansas Medical Center.

---

**Supplementary Table 1.**
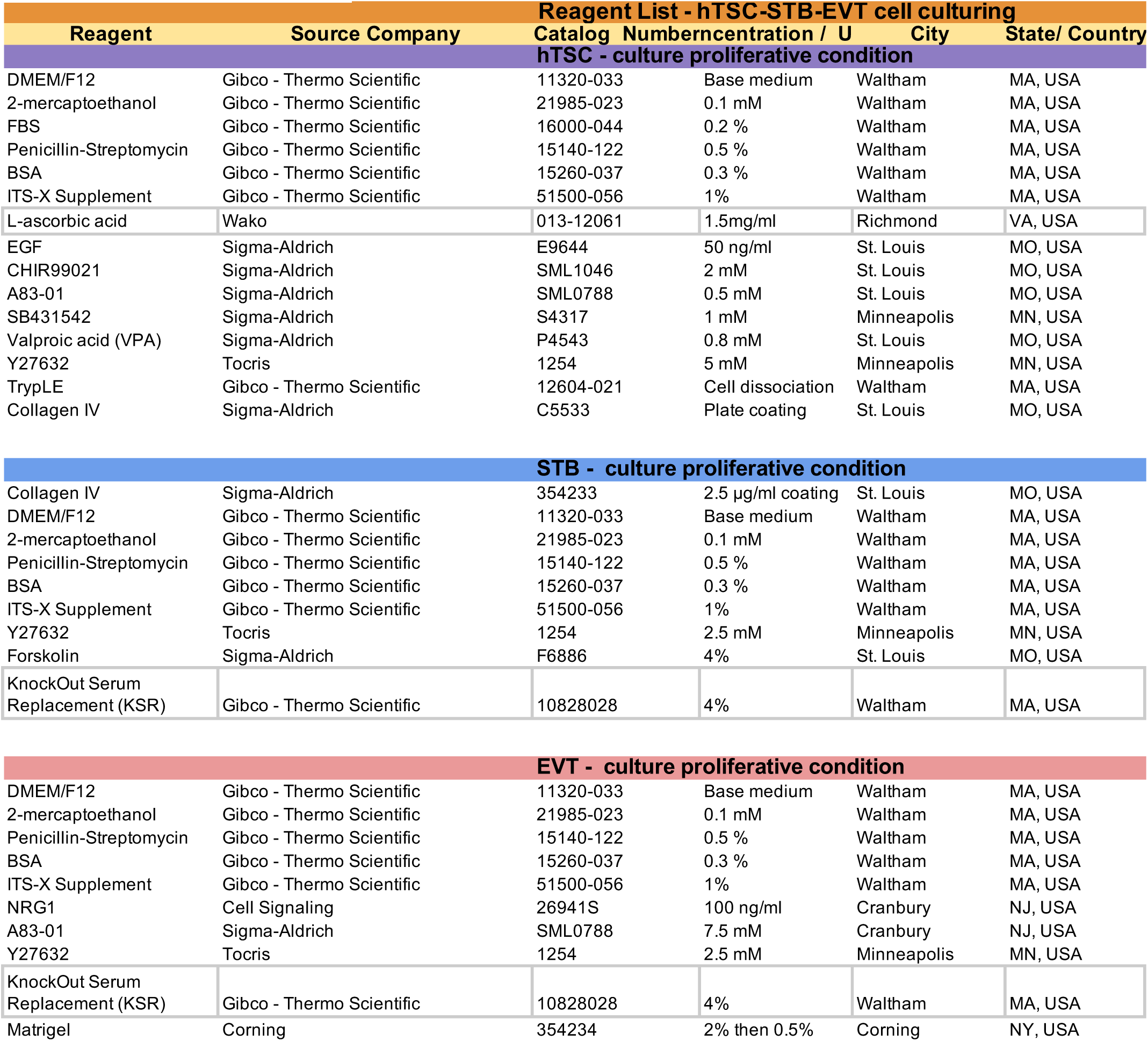

**Supplementary Figure S1:**
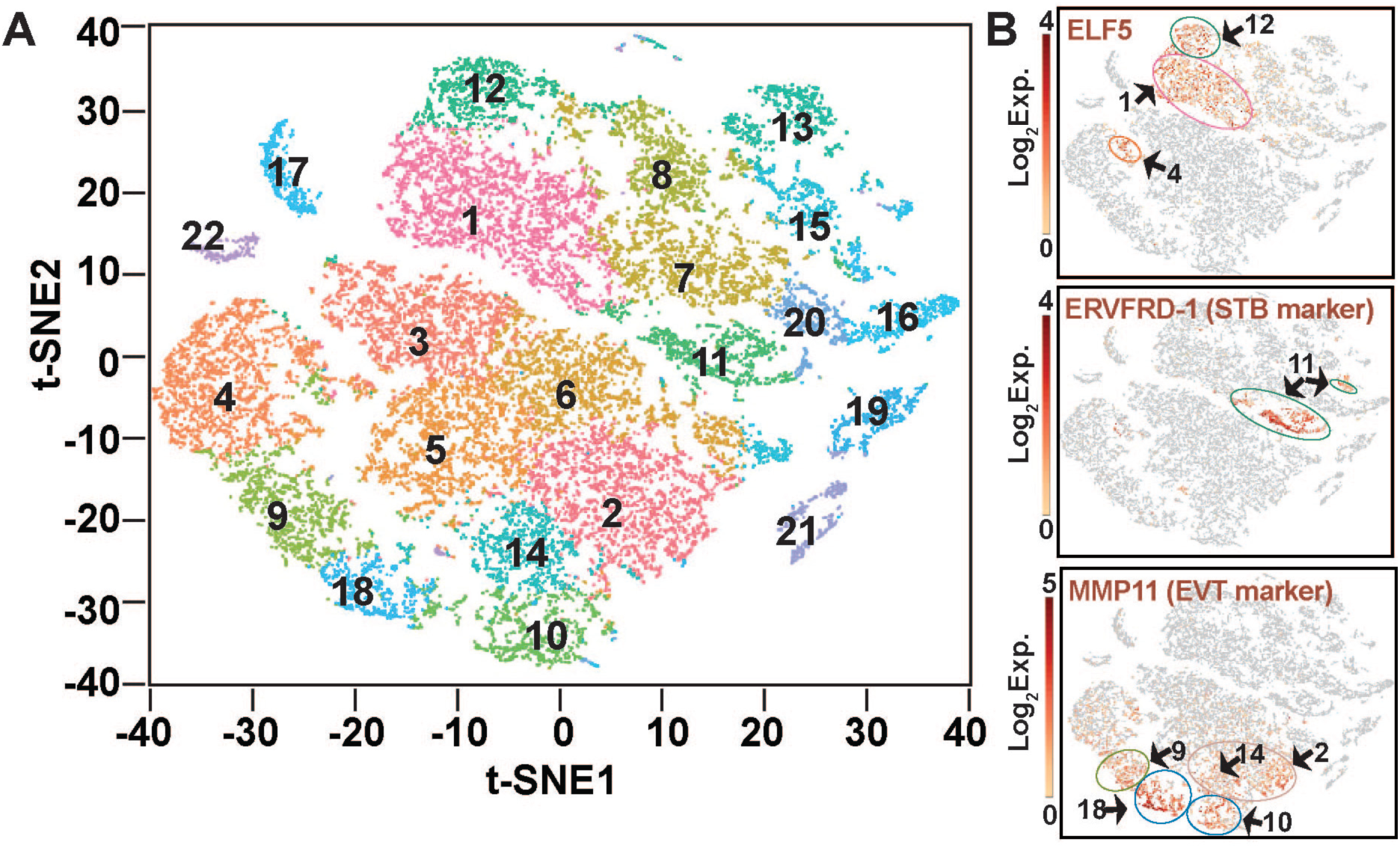
scRNA-seq analyses in first-trimester human placentae. (A) The t-SNE plot of hierarchical clustering of single cell RNA seq of human placental samples identified 22 different cell clusters. (B) Clusters of undifferentiated CTBs, STB-commutted CTBs and EVT precursors are identified from high expressions of ELFS, ERVFRD-1 and MMP11, respectively.

**Supplementary Figure S2:**
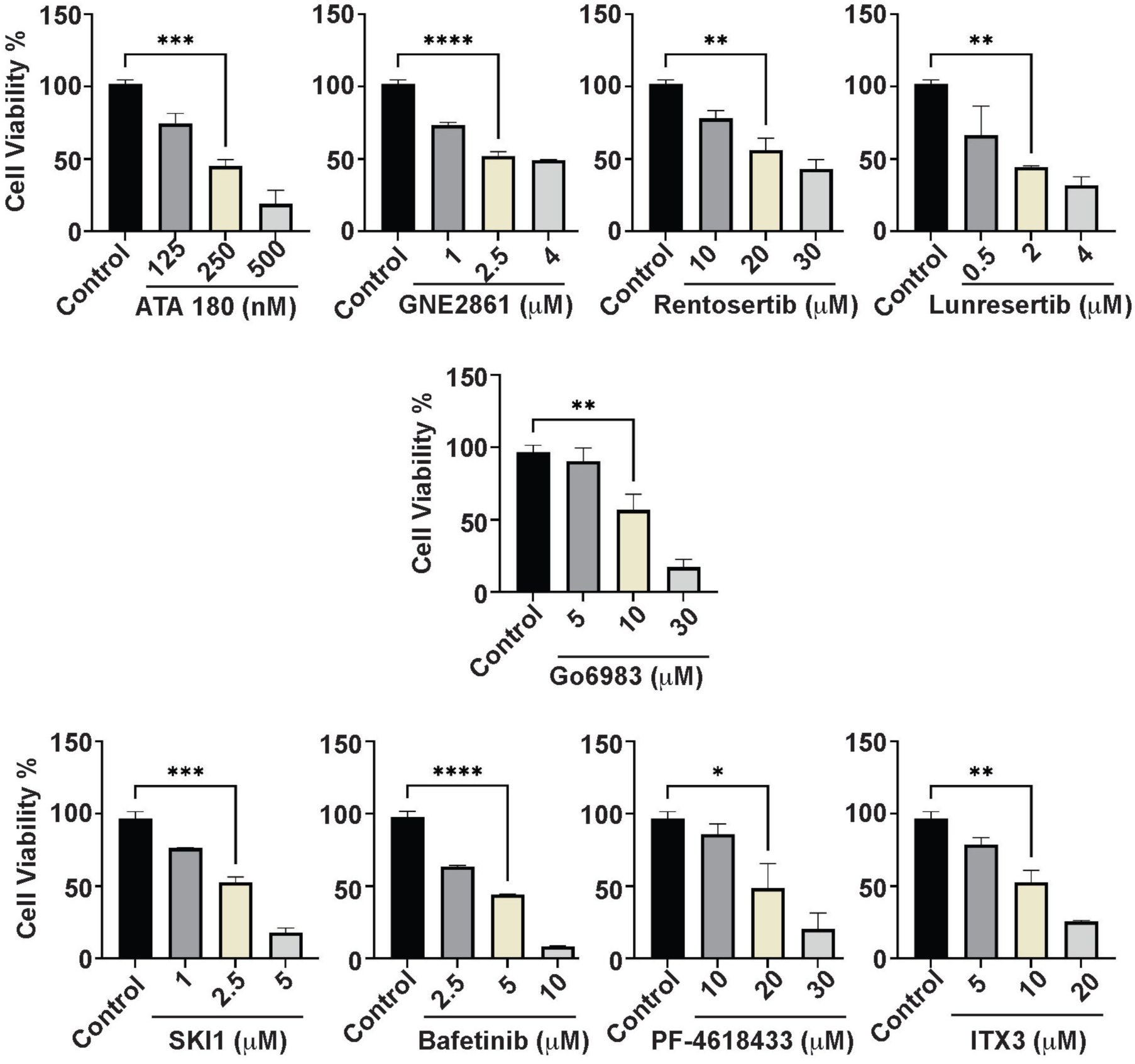
The Plots show viability of hTSCs, determine by MTT assay, when cultured with different concentrations of various kinase inhibitors.

**Supplementary Figure S3:**
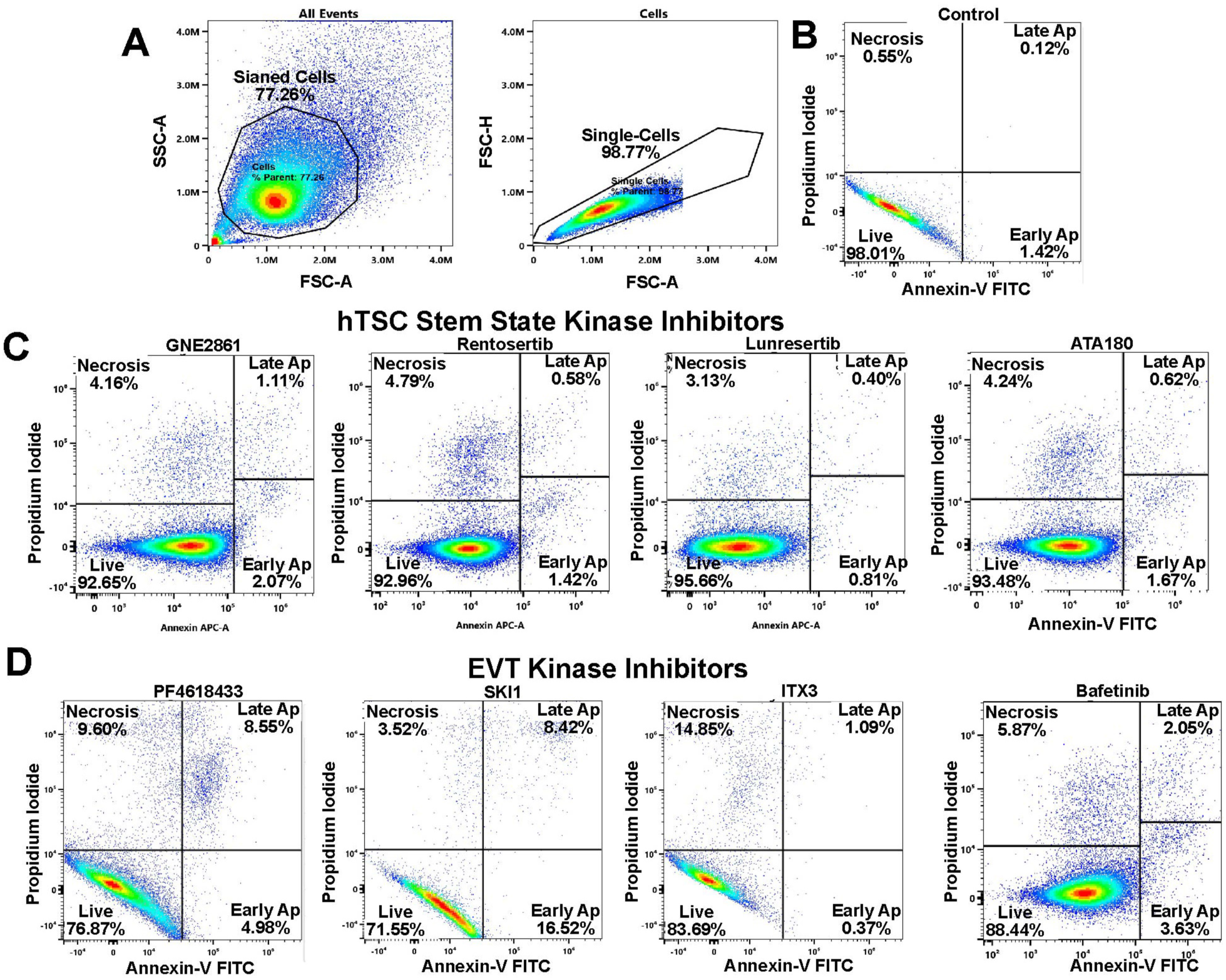
(A) Gating strategy to identify stained single cells. (B) Plot shows percentage of live and appoptotic cells when hTSCs were cultured for 3 days in stem-state culture condition with DMSO control. (C) and (D) Plots show percentage of live and appoptotic cells when hTSCs were cultured for 3 days in stem-state culture condition-with inhibitors of hTSC stem-state specific or EVT-specific kinases.

**Supplementary Figure S4:**
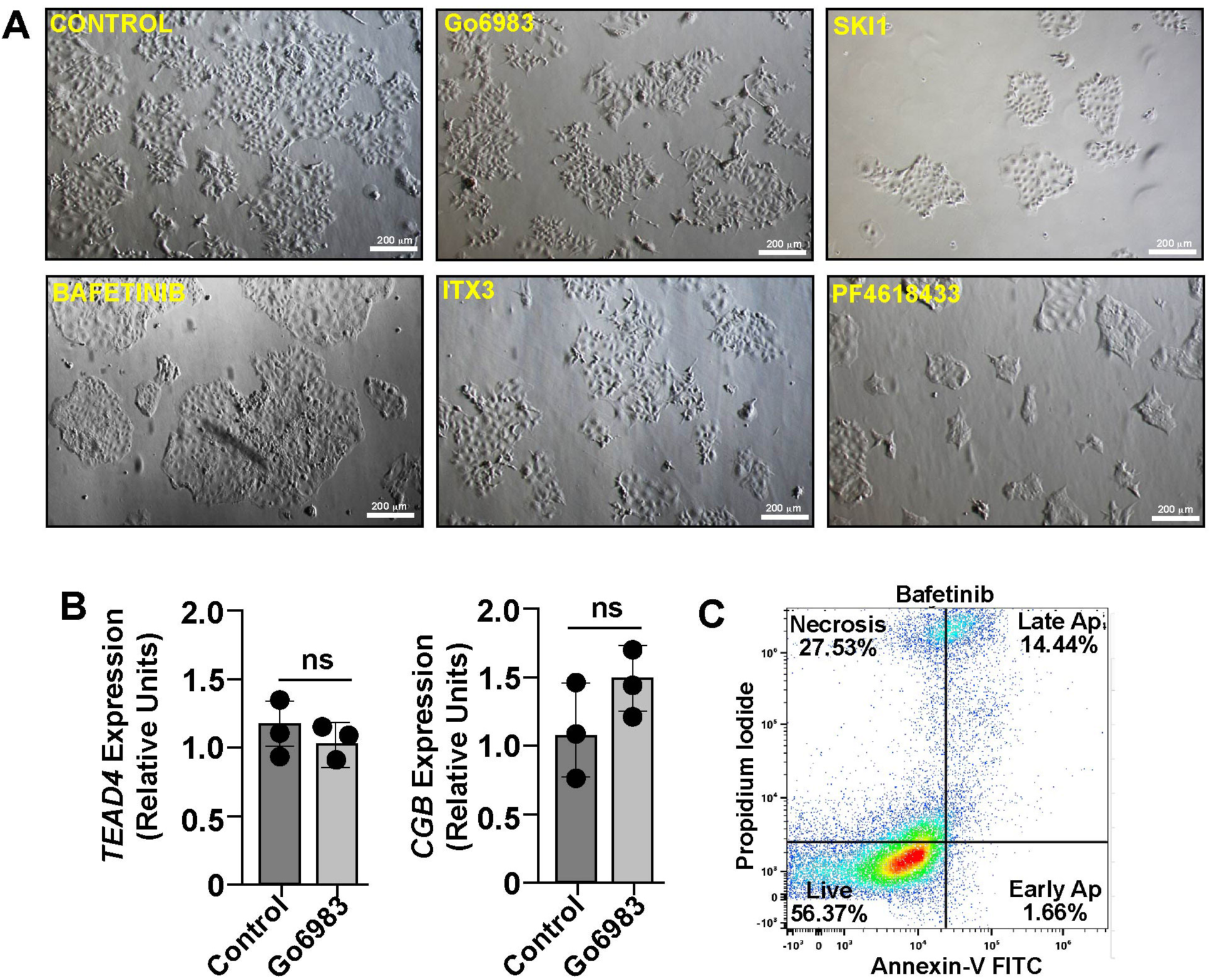
(A) Micrographs show human TSC colony morphologies, when cultured in stem state culture condition in presence of different kinase inhibitors, which impair either STB differentiation (Go6983) or EVT development (SKI1, Bafetinib, ITX3 and PF-4618433). Note that in presence of inhibitors cell death is not noticeable. However, partial reduction of TSC cell proliferation is apparent in presence of SKl1 and PF4618433. (B) RT-qPCR analyses of *TEAD4* and CGB expressions when hTSCs were treated with Go6983 in stem-state culture. (C) Plot shows percentage of live and appoptotic cells when hTSCs were cultured with Bafetinib in EVT culture condition.

