## Supplemental Tables and Figures for "An Integrated Proteomics and Genomics Approach to Identify Essential Protein Kinases During Human Trophoblast Development"

**Supplementary Table 1**

| Reagent | Source Company | Catalog Numberncentration / U | | City | State/ Country |
|---|---|---|---|---|---|
| **Reagent List - hTSC-STB-EVT cell culturing** | | | | | |
| **hTSC - culture proliferative condition** | | | | | |
| DMEM/F12 | Gibco - Thermo Scientific | 11320-033 | Base medium | Waltham | MA, USA |
| 2-mercaptoethanol | Gibco - Thermo Scientific | 21985-023 | 0.1 mM | Waltham | MA, USA |
| FBS | Gibco - Thermo Scientific | 16000-044 | 0.2 % | Waltham | MA, USA |
| Penicillin-Streptomycin | Gibco - Thermo Scientific | 15140-122 | 0.5 % | Waltham | MA, USA |
| BSA | Gibco - Thermo Scientific | 15260-037 | 0.3 % | Waltham | MA, USA |
| ITS-X Supplement | Gibco - Thermo Scientific | 51500-056 | 1% | Waltham | MA, USA |
| L-ascorbic acid | Wako | 013-12061 | 1.5mg/ml | Richmond | VA, USA |
| EGF | Sigma-Aldrich | E9644 | 50 ng/ml | St. Louis | MO, USA |
| CHIR99021 | Sigma-Aldrich | SML1046 | 2 mM | St. Louis | MO, USA |
| A83-01 | Sigma-Aldrich | SML0788 | 0.5 mM | St. Louis | MO, USA |
| SB431542 | Sigma-Aldrich | S4317 | 1 mM | Minneapolis | MN, USA |
| Valproic acid (VPA) | Sigma-Aldrich | P4543 | 0.8 mM | St. Louis | MO, USA |
| Y27632 | Tocris | 1254 | 5 mM | Minneapolis | MN, USA |
| TrypLE | Gibco - Thermo Scientific | 12604-021 | Cell dissociation | Waltham | MA, USA |
| Collagen IV | Sigma-Aldrich | C5533 | Plate coating | St. Louis | MO, USA |
| **STB - culture proliferative condition** | | | | | |
| Collagen IV | Sigma-Aldrich | 354233 | 2.5 µg/ml coating | St. Louis | MO, USA |
| DMEM/F12 | Gibco - Thermo Scientific | 11320-033 | Base medium | Waltham | MA, USA |
| 2-mercaptoethanol | Gibco - Thermo Scientific | 21985-023 | 0.1 mM | Waltham | MA, USA |
| Penicillin-Streptomycin | Gibco - Thermo Scientific | 15140-122 | 0.5 % | Waltham | MA, USA |
| BSA | Gibco - Thermo Scientific | 15260-037 | 0.3 % | Waltham | MA, USA |
| ITS-X Supplement | Gibco - Thermo Scientific | 51500-056 | 1% | Waltham | MA, USA |
| Y27632 | Tocris | 1254 | 2.5 mM | Minneapolis | MN, USA |
| Forskolin | Sigma-Aldrich | F6886 | 4% | St. Louis | MO, USA |
| KnockOut Serum Replacement (KSR) | Gibco - Thermo Scientific | 10828028 | 4% | Waltham | MA, USA |
| **EVT - culture proliferative condition** | | | | | |
| DMEM/F12 | Gibco - Thermo Scientific | 11320-033 | Base medium | Waltham | MA, USA |
| 2-mercaptoethanol | Gibco - Thermo Scientific | 21985-023 | 0.1 mM | Waltham | MA, USA |
| Penicillin-Streptomycin | Gibco - Thermo Scientific | 15140-122 | 0.5 % | Waltham | MA, USA |
| BSA | Gibco - Thermo Scientific | 15260-037 | 0.3 % | Waltham | MA, USA |
| ITS-X Supplement | Gibco - Thermo Scientific | 51500-056 | 1% | Waltham | MA, USA |
| NRG1 | Cell Signaling | 26941S | 100 ng/ml | Cranbury | NJ, USA |
| A83-01 | Sigma-Aldrich | SML0788 | 7.5 mM | Cranbury | NJ, USA |
| Y27632 | Tocris | 1254 | 2.5 mM | Minneapolis | MN, USA |
| KnockOut Serum Replacement (KSR) | Gibco - Thermo Scientific | 10828028 | 4% | Waltham | MA, USA |
| Matrigel | Corning | 354234 | 2% then 0.5% | Corning | NY, USA |

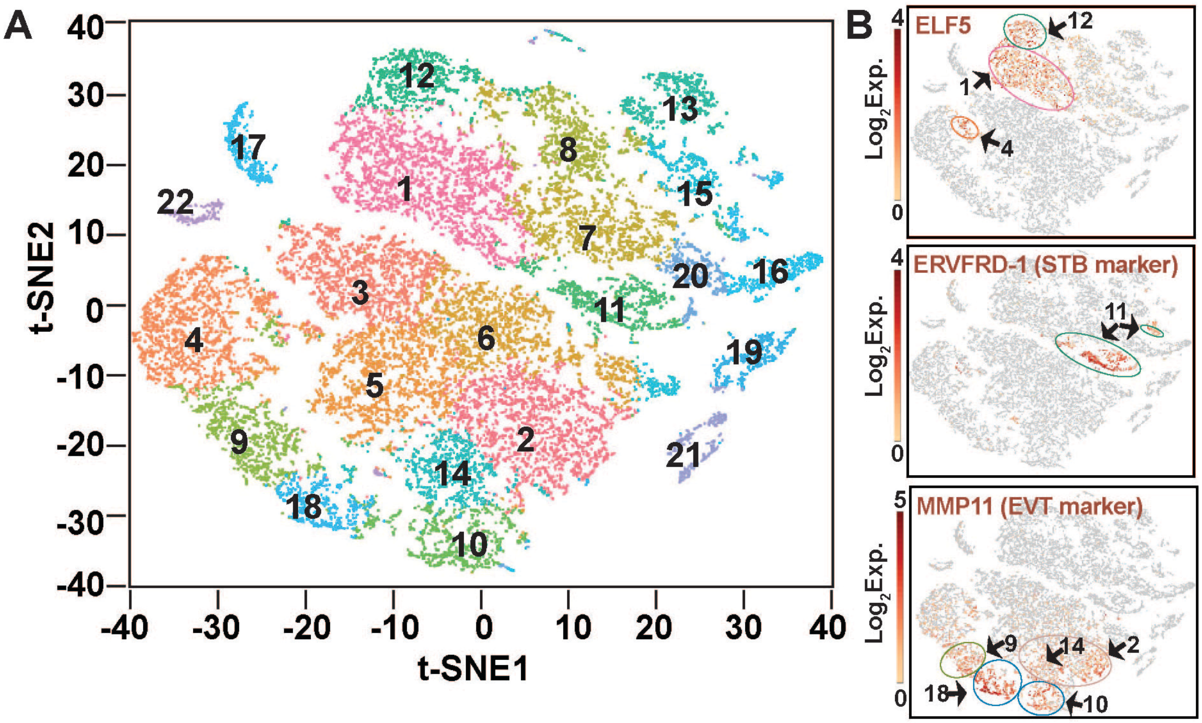


**Supplementary Figure S1: scRNA-seq analyses in first-trimester human placentae.** (A) The t-SNE plot of hierarchical clustering of single cell RNA seq of human placental samples identified 22 different cell clusters. (B) Clusters of undifferentiated CTBs, STB-commutted CTBs and EVT precursors are identified from high expressions of ELF5, ERVFRD-1 and MMP11, respectively.

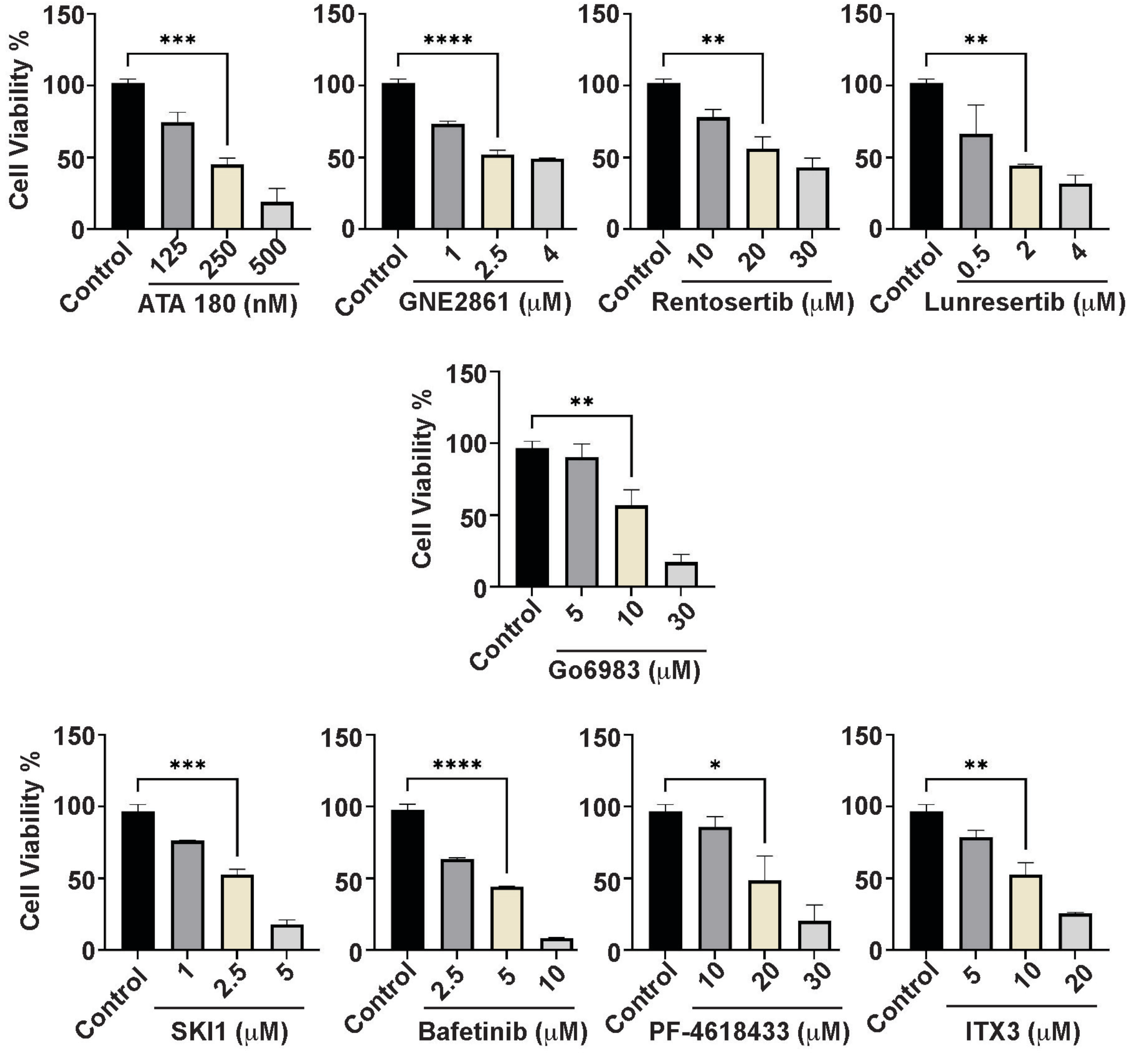


**Supplementary Figure S2:** The Plots show viability of hTSCs, determine by MTT assay, when cultured with different concentrations of various kinase inhibitors.

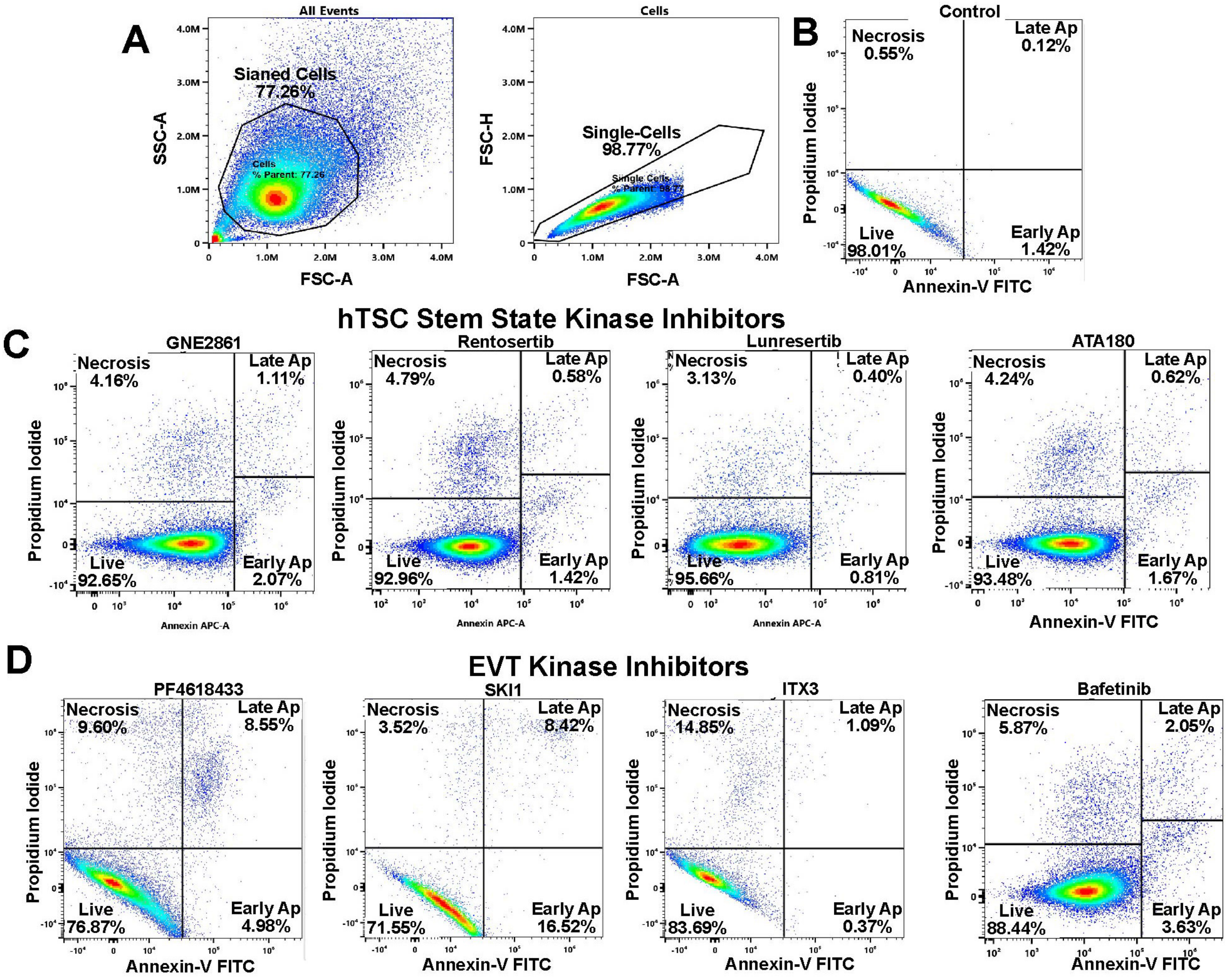


**Supplementary Figure S3:** (A) Gating strategy to identify stained single cells. (B) Plot shows percentage of live and appoptotic cells when hTSCs were cultured for 3 days in stem-state culture condition with DMSO control. (C) and (D) Plots show percentage of live and appoptotic cells when hTSCs were cultured for 3 days in stem-state culture condition-with inhibitors of hTSC stem-state specific or EVT-specific kinases.

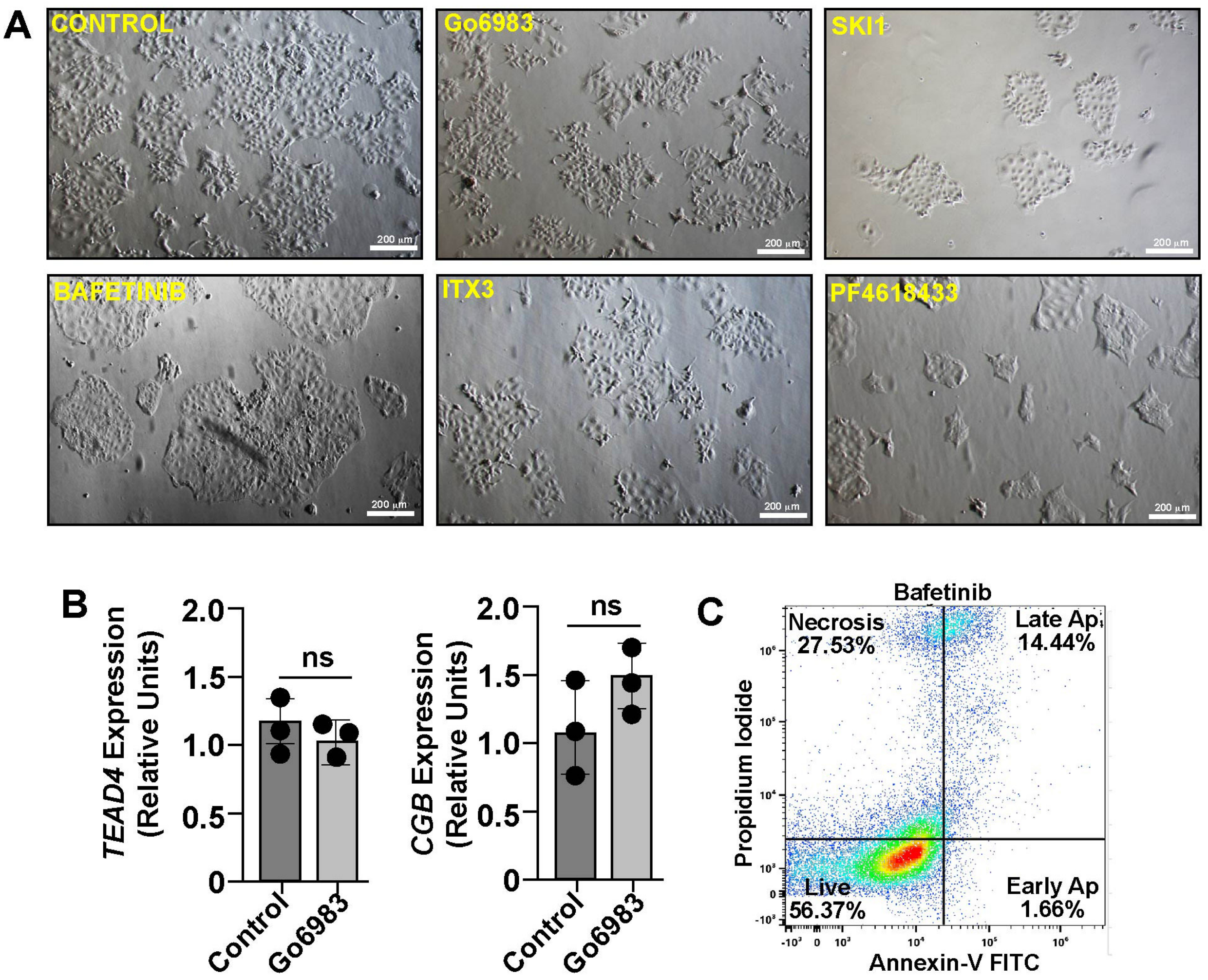


**Supplementary Figure S4:** (A) Micrographs show human TSC colony morphologies, when cultured in stem state culture condition in presence of different kinase inhibitors, which impair either STB differentiation (Go6983) or EVT development (SKI1, Bafetinib, ITX3 and PF-4618433). Note that in presence of inhibitors cell death is not noticeable. However, partial reduction of TSC cell proliferation is apparent in presence of SKI1 and PF4618433. (B) RT-qPCR analyses of *TEAD4* and *CGB* expressions when hTSCs were treated with Go6983 in stem-state culture. (C) Plot shows percentage of live and appoptotic cells when hTSCs were cultured with Bafetinib in EVT culture condition.
